# Zic3 enables bimodal regulation of tyrosine hydroxylase expression in dopaminergic neurons of olfactory bulb and midbrain

**DOI:** 10.1101/2022.10.30.514394

**Authors:** Smitha Bhaskar, Jeevan Gowda, Akshay Hegde, Surya Chandra Rao Thumu, Narendrakumar Ramanan, Jyothi Prasanna, Anujith Kumar

## Abstract

Dopaminergic (DA) neurons in the Olfactory bulb (OB) are involved in odor detection and discrimination. Transcription factor (TF) regulatory network responsible for their fate specification remains poorly understood and the spatial regulation of DA neurons remains elusive. In this study, mice exposed to odor stimulant exhibited specific upregulation of Zinc finger transcription factor of Cerebellum (ZIC) 3 along with Tyrosine Hydroxylase (TH). Stringent co-expression analysis showed ZIC3 and TH dual positive neurons in OB. Genetic manipulation showed ZIC3 to be both essential and sufficient to drive TH expression and essential for odor perception. ZIC3 interacts with ER81 and binds to region encompassing ER81 binding site in DA neurons and is indispensable for TH expression. In midbrain (MB), in the absence of ER81, ZIC3 switches its molecular partner and binds to *Pitx3* promoter-a DA fate determinant. Under ectopic expression of ER81 in MB DA neurons, propensity of ZIC3 binding to *Pitx3* promoter is compromised and its occupancy on *Th* promoter encompassing ER81 binding site is established, finally culminating in desired TH expression. Together, these findings reveal a unique ZIC3 mediated bimodal regulation of TH in OB and MB to ultimately facilitate DAergic fate.

## Introduction

Modulation in dopaminergic (DA) neuronal activity at different regions of the brain results in different disease phenotypes. Olfactory loss is one of the key non-motor symptoms (Xiao Q., et.al 2014) of Parkinson’s disease and serves as a faithful diagnostic marker (Morley et.al., 2018). Hyposmia-the reduced sense of smell manifests years before the first clinical symptom and is preceded by α-SYNUCLEIN accumulation in anterior olfactory nucleus (AON) (Rey et.al., 2018). One of the major reasons for hyposmia in PD patients is the predominant increase in tyrosine hydroxylase (TH) positive dopaminergic (DAergic) interneurons in the periglomerular layer (PGL) of Olfactory Bulb (OB) leading to heightened release of dopamine and consequent inhibition of olfactory signal transmission (Huisman et.al., 2004).

Neuronal diversity in OB is generated by fairly defined group of molecular determinants in a spatio-temporal manner, thus ensuring neurochemical and morphological variations in them (Parrish-Angust S., et.al 2007; De Marchis S., et.al 2007; Lledo P.M., et.al 2008). The complex developmental programs determining the OB DA neurons are orchestrated by transcription factors (TFs) PAX6 (Kohwi M., et.al 2005), ID2 (Havrda M., et.al 2005), ER81(Flames N., et.al 2009; Cave J.W., et.al 2010), MEIS2 (Agoston Z., et.al 2014), NGF1β (Saucedo-Cardenas O., et.al 1996), DLX2 (Brill M.S., et.al 2008), NEUROD1(Boutin C., et.al 2010) and PBX1(Remesal L., et.al 2020). In neonates, 4% of OB TH-positive cells are derived from EMX1-expressing progenitors, but in adult 42% of TH-positive OB interneurons are derived EMX1 expressing progenitors located in the dorsal portion of the SVZ (Kohwi M., et.al 2007). The TF DLX2 in a cell autonomous manner promotes the specification of DA neurons in OB (Brill M.S., et.al 2008). Over-expression of PAX6 in neuronal precursor cells enhanced the generation of DA interneurons and its association with MEIS2 is demonstrated to be a defining feature for DA fate specification (Agoston Z., et.al 2014). Another TF NEUROD1 plays a key role in terminal differentiation of OB DA neurons (Boutin C., et.al 2010) and so does the homeodomain TF PBX1(Remesal L., et.al 2020). Despite the discoveries on the role of some of these TFs, our understanding on the complex regulatory processes involved in the fine tuning of DAergic neurogenesis is still fragmentary.

Recent studies have been appreciating the pivotal role of TF ZIC3 (Zinc finger transcription factor in cerebellum 3), in neural development. ZIC3 belongs to ZIC family which comprises of ZIC1-5 members (Aruga J., et.al 1996; Nakata K., et.al 2000), and has domains similar to those of Gli proteins (Koyabu Y., et.al 2001). ZIC3 gene is present on X-chromosome and is highly conserved between mouse and human (Gebbia M., et.al 1997; Herman G.E., et.al 2022). Expression of ZIC3 is detected at embryonic two cell stage and later in adult animal the expression gets restricted to brain (Aruga J., et.al 1996; Kitaguchi K., et.al 2002). ZIC3 is highly expressed in ESCs and has been shown to be essential for maintenance of pluripotency and self-renewal of ESCs (Lim L.S., et.al 2007). Our previous study showed ZIC3 to assist efficient derivation of iPSCs from mouse fibroblasts in combination with OSK factors (Declercq J., et.al 2013) and further the mechanism was addressed by Sone et al., wherein it was showed ZIC3 along with ESRRB to synergistically enhance the glycolysis process and inhibit mitochondrial oxidative phosphorylation during the course of reprogramming (Sone M., et.al 2017). ZIC3 has also been demonstrated to regulate the transition of naïve to primed pluripotent state of pluripotent stem cells (Yang S.H., et.al 2019). In humans, the importance of ZIC3 is well reflected by its mutation leading to multiple disorders like heterotaxy (situs ambiguous) syndrome (Kitaguchi T., et.al 2000; Fritz B., et.al 2005), cerebellar anomaly (Purandare S.M., et.al 2002; Carrel T., et.al 2001) and anorectal malformation (Fritz B., et.al 2005).

ZIC family members are also known to play crucial role in neuronal development (Aruga J., et.al 1994). ZIC family members are some of the earliest TFs to be expressed during gastrulation and expressed earlier than that of proneural genes in the Neuroectoderm (Nakata K., et.al 2007; Mizuseki K., et.al 1998). BMP activity mediated expression of *zic1-3* in dorsal neuroectodermal marks the earliest event in neural fate determination (Marchal L., et.al 2009). Mice homozygous null for ZIC1 and ZIC3 have implications on expansion of Tyrosine hydroxylase (TH) positive neurons and caused hypoplasia of the hippocampus, septum, and OB (Inoue T., et.al 2007). ZIC3 knockout mice exhibits the bent tail phenotype featuring the neural tube defects (Klootwijk R., et.al 2000; Carrel T., et.al 2000). ZIC3 is also known to be involved in regulating the neural crest migration and differentiation by modulating the expression of neural crest genes FOXD3 and PAX3A (Garnett A.T., 2012; Sasai N., et.al 2001). A detailed expression profiling of ZIC3 in brain regions in the recent years has reported its presence in septal region and dorsal SVZ (Qin S., et.al 2017) thus suggesting its contribution to OB interneuron generation. Despite these insights, the consequences of ZIC3 absence on DA neurons generation or their maintenance are unclear.

In the present study, we report ZIC3 to be positively modulated upon exposing the animals to odor enriched environment and ZIC3 is expressed in a distinct micro-domain of OB DA neurons where it positively regulates TH expression. Interestingly, ZIC3 deploys distinct mechanism in OB and midbrain (MB) to fine tune the expression of TH and determine the DA fate. In the absence of a direct binding site on *Th* promoter, ZIC3 collaborates with ER81 in OB to drive the expression of *Th*, whereas in MB, ZIC3 indirectly dictates *Th* expression by occupying *Pitx3* promoter. Thus, we propose ZIC3 to regulate *Th* expression in a bimodal fashion and consequently DA fate in a spatially distinct manner.

## Materials and methods

### Animals

All animal experiments were performed in compliance with the guidelines of Association for the assessment and accreditation of laboratory animal care.

### Cell line culture

HEK293T cells were grown in DMEM High glucose medium supplemented with 10% FBS, 1X NEAA and 1X PenStrep. The cells were cultured till 80% confluence, split in the ratio of 1:5. mESCs were cultured on a layer of inactivated MEFs in medium containing DMEM high glucose-83.3%, FBS-16%, L-glutamine 2mM, anti anti-1%, NEAA-1X, sodium pyruvate-1%, β-ME-1.2% and supplemented with 8 ng/mL of Leukemia inhibitory factor (LIF). Prior to differentiation, mESCs were plated onto gelatin coated surface and grown for 2 days in medium containing LIF.

### Olfactory bulb DA neuron differentiation

Mouse embryonic stem cells (mESCs) were initially cultured on inactivated MEF feeder layer for at least 3 passages and later seeded on to 0.1% gelatin coated tissue culture dishes. After two days, the cells were trypsinzed using 0.05% trypsin and plated on to ultra low attachment dishes in ESC medium devoid of LIF to generate embryoid bodies (EBs) for 4 days. Later, the EBs were gently dissociated and plated at a density of 1 million cells/well of a 6 well plate with gelatin coating and allowed to attach in LIF deprived medium. Neural patterning was induced by culturing them in N2 medium supplemented with 2.5 ng/mL fibronectin and ITS. Medium change was provided every alternate day. Post 6^th^ day, the cells were grown in medium containing N2 supplement, bFGF, FGF-8 and purmorphamine for 4 days. Finally, the cytokines were withdrawn and the cells were allowed to achieve terminal maturation for 6 days in basal N2 medium.

### Generation of OB and MB neurospheres

E13-E15 mouse embryos were obtained from CCAMP and were subjected to cervical dislocation. The embryos were dissected using sterile scalpel blade to separate olfactory bulb (OB) and midbrain (MB) regions. The tissues were thoroughly washed with sterile 1X PBS supplemented with 0.5X PenStrep. OB and MB tissues were minced into fine pieces separately and dissociated using 0.25% trypsin for 10 minutes at 37°C. Trypsin was neutralized using 1:4 serum containing medium and the cells were spun at 1400 rpm for 5 minutes at room temperature (RT). After discarding the supernatant, the pellet was resuspended in neurospheres medium comprised of 1:1 DMEM F12 with 1X N2 supplement and Neurobasal medium with 1X B27 supplement with 2% FBS, 100 mM Glutamax, 100 µM β-Mercaptoethanol. The cells were plated onto ultra-low attachment plates for neurosphere formation. Medium change was performed every third day till day 12.

### Primary OB and MB DA neuron differentiation

To achieve dopaminergic (DA) neuron differentiation, OB or MB neurospheres were gently dissociated and plated onto 0.1% gelatin coated dishes at a density of 10*10^5^ cells/well of a 6 well plate in neurospheres medium. The plated cells were cultured as neural progenitors for 4 days *in vitro* (4 DIV) and further differentiation was induced by providing medium containing 1:1 DMEM F12 and Neurobasal medium supplemented with 1X N2, 100 mM Glutamax, 100µM β-Mercaptoethanol, 20 ng/mL bFGF, 100 ng/mL FGF-8 and 3 µM Purmorphamine. Differentiation was continued for 4 days *in vitro* (8 DIV) with alternate day medium change. Cells were fixed at 4 DIV and 8 DIV and characterized for the presence of appropriate markers to assess the efficiency of differentiation.

All cells were cultured in a humidified incubator (Thermo Fisher, USA) at 37°C and 5% CO2.

### Transduction

For the delivery of shRNA constructs, we utilized lentiviral packaging. HEK 293Ts were transfected using lentiviral packaging vectors Ps Pax2 and PMD 2G and the plasmid of interest. pLKO puro vector was used as the scrambled control for *Zic3* shRNA. The medium containing the lentiviral particles was collected at 48 hours and 72 hours post transfection. The viruses were concentrated by incubating with 50% PEG 6000 solution for 24 hours at 4°C on a rocker. The medium was spun at 1600 rpm for 1 hour at 4°C and the pelleted viral particles were used for transducing the recipient cells.

### Transcript Analysis: RNA isolation, cDNA synthesis and PCR

RNA was isolated from cultured cells by phenol-chloroform method using Trizol. The cells were washed with 1X PBS after decanting the medium and resuspended in Trizol. The RNA was isolated and stored at -80°C until use.

The isolated RNA was used to synthesize cDNA using commercially available first strand cDNA synthesis kit (# PGK162-A). The obtained cDNA was diluted 1:5 times using nuclease free water and further used for polymerase chain reaction.

Conventional PCR was performed by using Takara Emerald PCR Master Mix and the amplicons were observed on a 1.7% agarose gel. Real time PCR was performed using SYBR Green mix (#RR820B) with low ROX as the reference dye in 7500RT PCR system (Applied Biosystems, USA). Expression of the housekeeping gene *Gapdh* was set as control and the relative fold change for the gene expression was calculated.

### Immunofluorescence

Cells were washed thrice with 1X PBS and were fixed using 4% PFA for 20 minutes at RT and later washed thrice with 1X PBS. Permeabilization was done using 0.5% Triton-X 100 prepared in PBS for 30 minutes, followed by blocking using 20% FBS in PBS for 20 minutes. All steps were carried out at RT. Appropriate primary antibodies diluted in PBS were added to cells and incubated overnight at 4°C. Cells were washed thrice with 1X PBS to remove any non-specific binding. Fluorescent tagged secondary antibodies were added to the cells at the concentration of 1:800 (diluted in PBS) for 1 hour 30 minutes at RT. The cells were then washed with PBS and were counterstained with DAPI (1:10000 dilution) for 1 minute and again washed with PBS. Cells were observed and imaged using Olympus 1×73 inverted microscope under 20X magnification. Images were tinted and further processed using ImageJ software.

### Fluorescence activated cell sorting (FACS)

Trypsinized cells were washed with 1X PBS and later fixed using 0.5% Paraformaldehyde (PFA). The samples were stored at 4°C till use. Permiabilization was performed with 1X BD Permwash solution in PBS for 30 minutes at RT on a rocking platform. The non specific sites were blocked using 20% FBS in 1X BD Permwash for 20 minutes at RT. Primary antibody was added in 1X BD Permwash at a dilution of 1:400 and incubated at 4°C on a rocker overnight. The cells were washed with 1X PBS thrice and secondary antibody was added in 1X BD Permwash at a dilution of 1:800 and incubated at RT for 1 hour in dark. The antibodies used have been listed in Supplementary table 2. Washes were performed with 1X PBS thrice, the cells were resuspended in PBS and the events were acquired on LSR II flow cytometer using 488 laser line or on FACS ARIAFusion using 561 nm laser.

### Magnetic activated cell sorting (MACS)

About 10* 10^6^ OB cells were resuspended in 70 μL of buffer containing 0.5% BSA in 1x PBS. Further, 10 μL of FcR blocking reagent was added to the cells and incubated for 10 minutes at 4°C. The antibody microbeads were added at a volume of 20 μL per tube, mixed thoroughly and incubated for 15 minutes at 4°C. The cells were washed with 2 mL of buffer and centrifuged at 300*g for 10 minutes and the pellet was resuspended in 500 μL buffer. Magnetic separation was performed by passing the samples through MS columns. The unlabeled cells were obtained in the flow through and the magnetically labeled cells were plunged into a fresh tube. The separated cell populations were further labeled with PSA-NCAM antibody and the procedure was repeated. Finally, the MACS separated cells were co-stained for TH and ZIC3 and were analyzed on a flow cytometer with single color and negative controls.

### Cloning

For *Zic3*-GST cloning, the 1401bp (*Zic3*) was amplified using mESC cDNA as a template and the amplicon was gel eluted using gel elution kit (Abzyme). The vector backbone AIRAP9-PSMD2-GST (Addgene no. 21799), was digested using *HinDIII* and *XhoI* to remove the PSMD2 CDS and the amplified *Zic3* gene was cloned using Infusion kit (Clontech, USA) as per manufacturer’s instruction.. To obtain various truncated versions of ZIC3, regions of CDS without one or more zinc finger domains were amplified and cloned between HinDIII and XhoI regions of GST tag vector. To clone ER81 downstream a 3X FLAG tag, FLAG-FUS WT vector backbone (Addgene no. 44985) was digested with *KpNI* and *XhoI* enzymes and the FUS CDS is replaced by ER81 CDS which was amplified using mouse OB cDNA.

### Western blot analysis

Cells were lysed using RIPA buffer after adding protease inhibitor cocktail, sodium orthovanadate and phosphatase inhibitors and were sonicated. The samples were spun at 12000g for 20 minutes at 4°C and the supernatant was collected. The isolated proteins were stored at -80°C until further use.

Proteins were fractionated on a 12% SDS PAGE and the resolved proteins were transferred onto an activated PVDF membrane for two hours at 100mA using a semi dry transfer apparatus. The blots were blocked using 5% skimmed milk and later incubated with respective primary antibodies at 4°C, overnight on a rocking platform. Three washes were performed using 1X TBST (pH 7.4) after which, HRP-tagged secondary antibodies were added at a final concentration of 1:2000 in skimmed milk and incubated at room temperature for 1 hour. After three washes with 1XTBST, the signal was developed using WesternBright ECL substrate (#K120445-D20) and captured in Li-COR chemiluminescence detector.

### Chromatin immunoprecipitation (ChIP)

To assess binding of ZIC3 to *Th* promoter, a ChIP assay was performed using Sigma Imprint chromatin immunoprecipitation kit (Sigma-CHIP1-24RXN) according to the manufacturer’s protocol. Briefly, about 4 μg of ZIC3 or ER81 antibody was allowed to bind to the assay wells washed with antibody buffer and incubated on a rocker with 120 rpm for about 90 minutes. Excess antibody was removed by performing 6 washes with IP wash buffer and a final wash with Tris-EDTA buffer.

The samples were prepared by making single cell suspension of the OB tissue derived from adult Swiss albino mouse. The cross linking was performed using 37% formaldehyde in serum free medium.. The nuclear components were obtained by resuspending the cells in nuclei preparation buffer. The obtained DNA was resuspended in shearing buffer with protease inhibitor cocktail (1X) and incubated on ice for 10 minutes. The sample was briefly vortexed and sonicated using 7 pulses thrice with an interval of 10 minutes between the cycles. The sheared chromatin was spun at 10000*g for 10 minutes at 4° C and the supernatant was used for ChIP further.

The lysate was diluted 1:1 with the dilution buffer and incubated in the antibody bound wells on a rocker at 120 rpm for 90 minutes. The wells were washed with IP wash buffer and the DNA release buffer containing proteinase K was added. The samples were incubated at 65° C for 15 minutes. Reverse cross linking was carried out for the samples by adding the buffer and incubating for 90 minutes at 65° C. The solutions were added onto a DNA binding column and purified to obtain chromatin and used for targeted realtime PCRs. A 5% input of the lysate was used as a control and the fold enrichment was plotted.

### Co-immunoprecipitation

OBs dissected from adult Swiss albino mouse were minced and trypsinized using 0.25% trypsin for 10 minutes at 37° C. The cell pellet was resuspended in IP lysis buffer containing Tris (20mM, pH-8.0), NaCl (137 mM), NP-40 (1%), EDTA (2 mM) and protease inhibitors and incubated on a rocker at 120 rpm for half an hour. The cell lysate was spun at 12000*g for 10 minutes at 4°C and the supernatant was collected. The supernatant was pre-cleared by incubating with protein-A sepharose bead slurry on a rocking platform for 4 hours at 4°C. The samples were spun at 4°C at 1200 rpm for 5 minutes and the supernatant was collected as the pre-cleared lysate. The Protein-A slurry was incubated with 1-4μg of the appropriate antibody on a rocking platform at 4°C overnight. Later, the pre-cleared lysate was added to this mixture and incubated at 4°C overnight on a rocker. Immunoprecipitation was performed by eluting the complex of sample antibody from the beads by inducing pH change with 0.2 M glycine (pH 2.0) followed by neutralization with 1M Tris (pH 7.0). The immunoprecipitated samples were then used for western blot analysis.

### Immunohistochemistry

OBs of 3 week old Swiss Albino mice were obtained and subjected to immunohistochemical staining using TH or ZIC3 antibody. The tissue was fixed with 4% PFA at 4°C for 72 hours and cryo-sectioning was performed to obtain 10 µm thick brain sections. The sections were washed once with 1X PBS and permeabilized with 0.5% triton X100. Blocking was performed using 20% normal Donkey serum and 1% BSA and the sections were incubated with 1:100 ZIC3 primary antibody (ab222124) overnight at 4°C. Washes were performed with 1X PBS and exposed to secondary antibody diluted in PBS to a final concentration of 1:600 and incubated for 2 hours at room temperature. The sections were washed thrice with 1X PBS and counter-stained with 1:10000 DAPI. The stained neurons were imaged under 20X objective in Nikon TE 2000 inverted epi-fluorescence microscope.

### Olfactory training to Mice

To determine ZIC3 expression pattern in mice with olfactory enrichment, 3–4-week-old Swiss Albino female mice were selected and categorized into control and test groups. Control animals were housed in a completely odor free environment and were prevented from exposure to any natural or artificial olfactory stimuli for 20 days. The test mice were directly exposed to various food-related olfactory stimuli for 1 hour every alternate day such that their environment is enriched with dopamine stimulating odor molecules. Both the groups were maintained on same diet and their bedding was changed every day. At the end of the olfactory training, animals were subjected to cervical dislocation and OBs were harvested for either transcript or protein analysis.

### Luciferase assay

Cells were grown to 80% confluence and transfected with 50 ng TK-Renilla and 950 ng of Th-luciferase plasmids. After 24 hours of transfection, medium was changed and the cells were allowed to grow for another one day. The cells were harvested with appropriate volume of passive lysis buffer and stored at -80°C until further use. Luciferase assay was performed according to manufacturer’s instruction using dual luciferase assay kit (Promega Corporation). Readings were obtained using Ensight multiplate reader.

### Stereotaxy

Custom made lentivirus carrying shRNA targeting *Zic3* was injected in 7 months old wild type BL-6 mice. Briefly, the mice were placed in anaesthesia chamber for induction with 2-3%isoflurane(Sosrane, NEON, Mumbai, India). Post induction, the mouse was placed on the stereotaxic apparatus (KOPF, TUNJUNG, CA, USA) with continuous supply of isoflurane of 1-1.5% for maintenance until the viral injections were completed. The injections were made in the substantia nigra pars compacta (SNpc) using 10 µl Hamilton syringe fitted with a 28-gauge needle (Hamilton Company, Nevada, USA) using the following stereotaxic coordinates in mm with reference to bregma: anteroposterior (AP) = 3.0, Mediolateral (ML) = 1.2, and dorsoventral (DV) = 4.5, with a flat skull position (Paxinos and Franklin). The viral injections were made at a speed of 0.1 μL per minute (1 μL, of 5× 10^12^ viral particles per mL) followed by an additional 5 min for complete diffusion of the virus and 3 min for slow withdrawal of the needle. Post injections the mouse was warmed under infrared (IR) lamp (245 V, 150 W) until being awakened and returned to the cage. The animals were maintained for 10 days and transcardially perfused. The brains were isolated, fixed in 4% paraformaldehyde (PFA) overnight, cryoprotected in 30% sucrose, frozen and sectioned at 30 µm in a cryostat (Leica, CM 1850).

### Statistical analysis

All the experiments (unless mentioned in the figure legend) were performed in biological triplicates with added technical replicates wherever applicable. The values were averaged, and standard errors were calculated. Student’s T tests were performed to determine the statistical significance and the graphs were prepared using GraphPad Prism 8.0. Significance was decided as follows: p≤ 0.05=*, p≤ 0.005=**,≤ 0.001=***

## Results

### Zic3 is associated with odor perception in OB DA neurons

Modulation of dopamine levels positively correlates with the odor perception or the sense of smell. Exposure of animals to natural odor stimulants induces dopamine pathway and ensures heightened olfactory discrimination and learning. The transcription factors directly involved in TH regulation respond to modulation in odor stimulation as seen in the case of NURR1 (Liu and Baker, 1999) and ER81(Cave et.al., 2010). In our study, upon olfactory enrichment with exposure to natural food related odorants, the animals exhibited an anticipated up-regulation in the transcript levels of *Th* by 3.5 fold and *Omp* by 9 fold along with increase in *Er81* expression (Figure 1a.i). As previous studies had reported a drastic decrease in the size of olfactory bulb in absence of *Zic1* and *Zic3* (Inoue et.al., 2007), we screened for the expression of *Zic* family members in olfactory bulb upon odor stimulation and found that while *Zic3* showed a significant up-regulation with respect to control, *Zic1* and *Zic2* failed to positively respond (Figure 1.a.ii). Protein analysis showed a significant ∼9% increase in TH expressing cells (Figure 1.b.i and ii) with a concomitant 12.5% increase in ZIC3 positive cells (Figure 1.b.iii and iv) in animals housed in an odor enriched environment in comparison with odor deprived control group. These results advocated ZIC3 as a probable mediator of dopamine production in response to odor enrichment.

**Figure 1:**
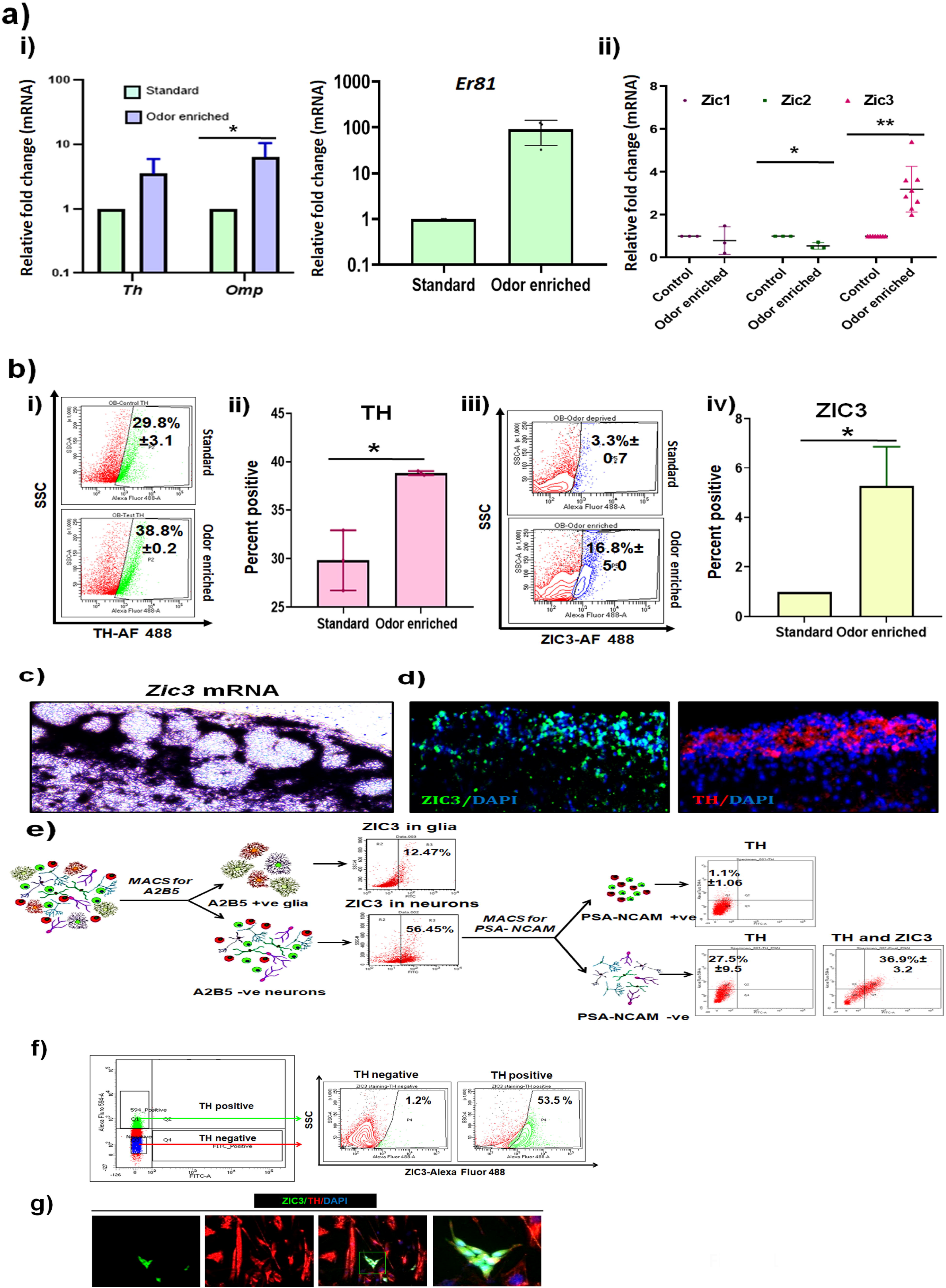
Zic3 is associated with odor perception in olfactory bulb DA neurons. (a) Transcript analysis of *Th, Omp, Er81* (i) and *Zic* family genes (ii) in mice exposed to odor enriched environment in comparison to control condition, (b) analysis of TH positive (i) and ZIC3 positive (ii) stained cells by flow cytometry, and quantification of flow cytometry analysis showing the fold change in expression levels of TH (ii) and ZIC3 (iv) in animals under odor enrichment, (c) in situ hybridization and (d) immunohistochemical staining for ZIC3 and TH representing their presence in the outermost periglomerular layer of mouse OB, (e) MACS to separate PSA-NCAM-ve and PSA-NCAM+ve cells from mouse OB and analysis for co-expression of TH and ZIC3, (f) FACS to fractionate TH+ve and TH-ve cells followed by flow cytometric analysis to score for co-expression of ZIC3 and TH, (g) Expression of TH in OB primary neurons harboring *Zic3*-promoter GFP. Mean+/-SE of biological triplicates, * p≤ 0.05.

Olfactory bulb (OB) is a highly complex structure organized into six main layers. Morphologically, neurochemically and functionally distinct neurons occupy different layers of OB forming the dopaminergic and CalR/CalB positive neurons in the outermost periglomerular layer (Bonzano et.al., 2016), glutamatergic neurons in the mitral and tufted layers and external pleaxiform layer (EPL) being mainly composed of dendritic synapses of mitral and tufted cells (Nagayama et.al., 2014). These distinct neurons are generated as a result of specific combinations of TFs. Mapping the expression domains of ZIC3 in mouse OB revealed that *Zic3* expression is confined to brain tissue and was undetectable in liver and heart, which represent the endodermal and mesodermal origin respectively (Figure S1a.i). However, within the mouse brain, we found *Zic3* distribution to be across cerebrum, cerebellum, OB and brain stem (Figure S1.a.ii). We confirmed this by flow cytometry analysis that showed the presence of ZIC3 positive cells in various brain regions (Figure S1.b.i and ii). The specific staining of brain cells and not the liver cells also authenticated the specificity of the ZIC3 antibody used in the study. Spatial distribution analysis of *Zic3* in OB by *in situ* hybridization showed the presence of cells expressing *Zic3* mRNA in the outermost periglomerular layer (Figure 1c). Immunohistochemical staining also demonstrated ZIC3 positive cells in the periglomerular layer of OB which is primarily constituted by the TH positive dopaminergic interneurons (Figure 1d). To find out if ZIC3 expression is enriched in neural stem/progenitor cells or the matured neurons of OB, we adopted Magnetic associated cells sorting (MACS) to segregate the OB neuronal population and determine ZIC3 expression. Initially, by labeling P2 mouse derived OB cells with A2B5 antibody, we separated A2B5 positive glial cells from A2B5 negative neuronal population. Flow cytometry analysis showed 56% of OB neuronal cells to express ZIC3 in comparison to 12% in glial cells (Figure 1e). Neuronal cells were further labeled with PSA-NCAM antibody to fractionate PSA-NCAM positive neural progenitors from PSA-NCAM negative matured neurons. While 1% of PSA-NCAM positive cells were TH positive, about 27.5%±9.5 of PSA-NCAM negative cells expressed TH. About 36.9%±3.2 of the PSA NCAM negative cells were both ZIC3 and TH positive, thus establishing the presence of a subset of OB DA neurons that harbor ZIC3. In an independent approach, we used FACS to segregate TH positive and negative DA neurons, wherein 20% of the cells were positively labeled for TH. Sequentially, fractionated TH positive and negative cells were stained with ZIC3 and analyzed for their co-expression. Among the sorted cells, about 32.3% of TH positive cells co-expressed ZIC3 (Figure 1f) reiterating the presence of ZIC3 in a subset of OB DA neurons. To corroborate this observation, we used E13.5 mouse OB primary neurons to transduce with *Zic3* promoter driven GFP plasmid. Post 3 days of transduction, the cells were fixed and stained using TH antibody. Detection of TH in ZIC3-GFP positive primary neurons (Figure1g) conclusively indicated the expression of ZIC3 in OB DA neurons. Thus, we report ZIC3 to be co-expressed in a subset of OB DA neurons.

### ZIC3 is essential for generation and maintenance of TH+ve cells of olfactory bulb

In a quest to understand if ZIC3 expression is essential to confer DAergic like identity, we chose mouse OB derived primary neurons obtained from E12.0-E14.0 mouse embryos as a model system. The OB identity is confirmed by scoring for the presence of mixed OB interneuron population represented by co-expression of GABA recB with TH, CalB, CalR or ER81 and few differentiated neurons expressing TUJ1. Absence of a midbrain DA specific marker PITX3 reassured the exclusive OB identity of these cells (Figure S2a). Upon culturing the OB progenitor cells as neurospheres, the cells expressed progenitor markers SOX1 and NESTIN (Figure S2b) and upon further culturing for 4 days in presence of FGF2, FGF8 and Purmorphamine the cells were differentiated to TUJ1 +ve (Figure S2b) and TH+ve DA like neurons with a co-expression of GABA rec1B-a hallmark of OB DA neurons (Figure S2c). Using this OB primary neuron model system, we performed genetic inhibition of *Zic3* by specific shRNA against 3’ UTR region. The efficiency of shRNA is confirmed at the transcript level by analyzing the inhibition of *Zic3* mRNA (Figure S3a) and at functional level by observing the down-regulation of NANOG protein in mESCs (Figure S3b). Down-regulation of *Zic3* in differentiated OB primary neurons failed to express *Th* in comparison to scrambled control alongside down-regulation of olfactory marker protein (*Omp*) (Figure S3c). To examine if the effect was indeed by the virtue of *Zic3* modulation, we performed a rescue experiment by reconstituting *Zic3* expression in OB primary cells harboring *Zic3* shRNA. The ZIC3 overexpression plasmid authenticity was confirmed (Figure S3.e) and observed to upregulate *Th* expression in OB primary cells when the cells were subjected DA neuronal differentiation(Figure S3.e). In the cells overexpressing *Zic3*, the over-expression restored the *Th* expression which was inhibited by *Zic3* shRNA(Figure 2a). Expression of GABA rec1B-remained unaltered in presence or absence of ZIC3 (Figure S3d). Immunostaining for TH (Figure 2b.i) revealed the cells with *Zic3* shRNA to down regulate TH expression by 28.9%±2.99 as compared to scrambled control, which was rescued upon *Zic3* over-expression (Figure2b.ii). Flow cytometry results (Figure 2c.i) further supported the immunostaining analysis wherein TH was expressed only in 76%±1.2 of the differentiated OB primary neurons with ZIC3 shRNA as compared to scrambled control with 92%±1.1 of the cells expressing TH protein. (Figure 2c.ii). As the members of ZIC family are highly conserved and share common promoter binding sites, we were compelled to investigate the specificity of above said observations. To this end, *Zic1* inhibition (Figure S3.f) was performed and we failed to detect any significant modulation in either the *Th* transcript (Figure S3.f) or protein expression (Figure S3.g), thus advocating a novel and specific role for *Zic3* in TH regulation. To understand whether the role of ZIC3 is restricted to influence the DA neurogenesis or is also involved in the maintenance of DA identity after differentiation, expression of *Zic3* was inhibited in terminally differentiated neurons and scored for TH expression. Inhibition of *Zic3* led to a drastic down-regulation of both *Th* and *Omp* mRNA (Figure2d.i) and TH protein (Figure 2d.ii). Thus, ZIC3 plausibly participates in maintenance of TH in DA neurons apart from influencing its initial expression. To conclusively ascertain the role of ZIC3 in DA neuron fate specification, we over-expressed ZIC3 in undifferentiated OB primary cells and cultured them without any differentiation cues alongside a vector control. Ectopic expression of ZIC3 in neural progenitor cells cultured without any differentiation cues was sufficient to trigger the expression of *Th*, both at mRNA (Figure 2e.i) and protein (Figure 2e.ii) levels, indicating ZIC3 is sufficient to drive the expression of TH in neural progenitor. With all these results, we infer that ZIC3 is essential and facilitates DA phenotype establishment *via* TH regulation.

**Figure 2:**
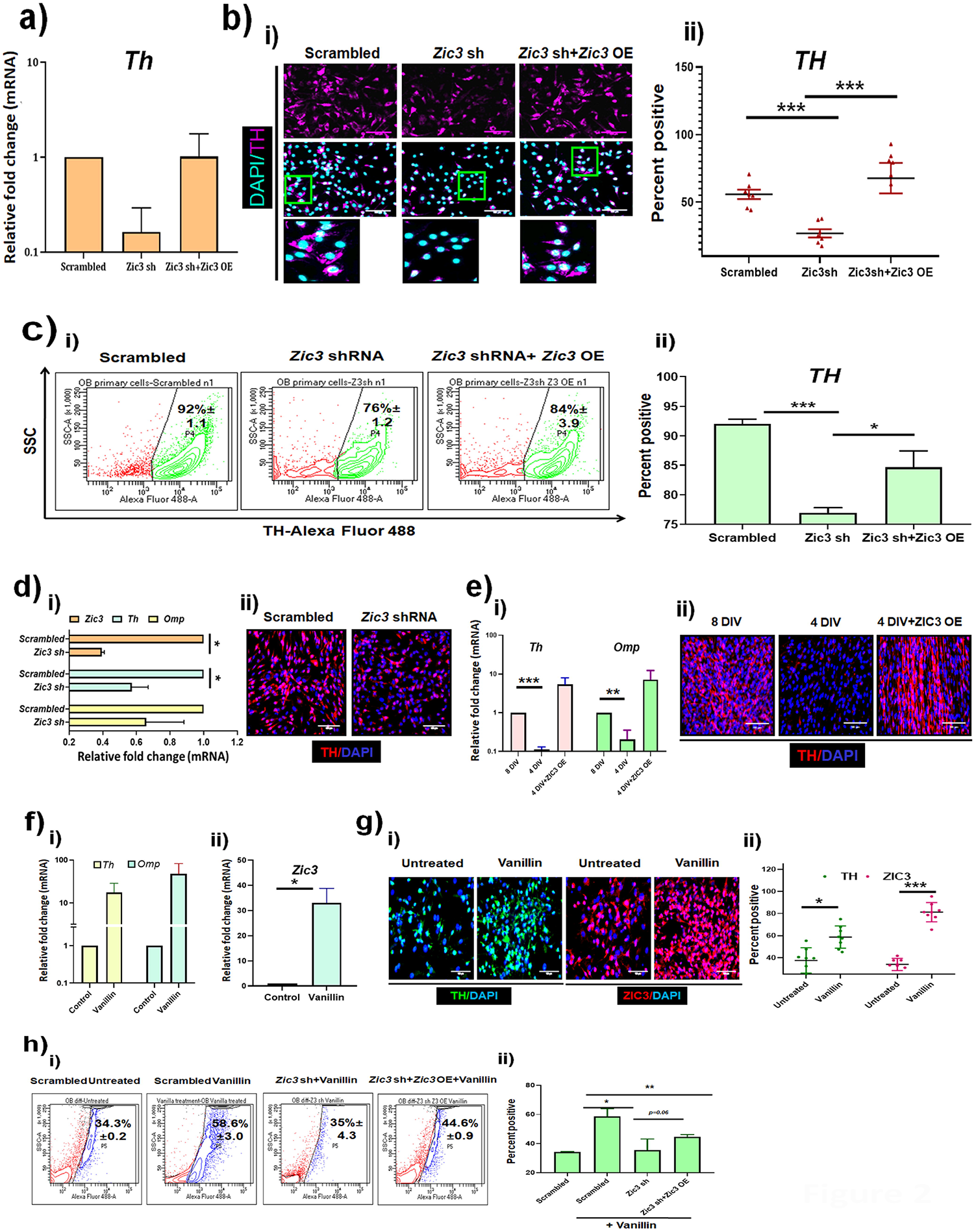
ZIC3 regulates TH expression in OB derived primary DA interneurons. (a) Transcript analysis of *Th* in OB DA primary neurons with *Zic3* shRNA and the rescue in expression by simultaneous overexpression of *Zic3*, (b) immunofluorescence staining for TH (i) and its quantification (ii) in OB DA like neurons derived in absence of *Zic3* and simultaneous overexpression of *Zic3* in shRNA condition, (c) expression of TH (i) and its quantification (ii) upon *Zic3* inhibition and over-expression as shown by flow cytometry, mRNA (d i) and protein expression (d ii) of TH in OB DA like neurons with *Zic3* inhibition post differentiation, (e) transcript (i) and protein (ii) analysis of TH expression in OB cells differentiated in presence and absence of differentiation cues and overexpression of ZIC3 in undifferentiated cells, (f) transcript analysis of *Th, Omp* (i) and *Zic3* (ii) in OB DA primary neurons with and without vanillin treatment, (g) immunofluorescence staining for TH and ZIC3 (i) and quantification graph (i) in cells treated with vanillin treatment and subjected to *Zic3* shRNA and overexpression conditions, (h) flow cytometry plots (i) and quantification (ii) of TH positive cells in scrambled, *Zic3* shRNA, *Zic3* sh+ *Zic3* OE conditions upon vanillin treatment. Scale bar represents 100 µm. Mean+/-SE of biological triplicates, * p≤ 0.05, ** p ≤ 0.01, *** p ≤ 0.001.

One of the major functions of DAergic neurons of OB is their involvement in sensing and relaying olfactory signals, mediated by TH expression. To determine if ZIC3 is necessary for up-regulation of TH in OB DA primary neurons in response to olfactory related stimulation, we treated the OB cells for 24 hours with Vanillin and performed transcript analysis. Cells treated with vanillin demonstrated a 47 fold increase in *Th*, 17 fold increase in *Omp* with respect to the untreated control (Figure 2f. i) along with approximately 70 fold induction in mRNA levels of *NeuroD1* (Figure S3.h) a TF involved in terminal maturation of OB DA neurons. A significant 33 fold increase in *Zic3* mRNA (Figure 2f. ii) was detected in vanillin treated cells with respect to the untreated control. Immunofluorescence imaging revealed a statistically significant increase in numbers of TH and ZIC3 positive cells (Figure 2g. i) in response to vanillin treatment which accounted for about 20% and a 43% respectively (Figure 2g. ii). While the scrambled control cells treated with vanillin were capable of increasing TH protein, knockdown of *Zic3* failed to maintain the TH expression (Figure 2h. i). However, TH expression was rescued in vanillin treated cells by 9.6% when *Zic3* was ectopically expressed in cells harboring *Zic3*sh RNA (Figure 2.h.ii). The specificity of *Zic3* in TH modulation is appreciated as another closely related ZIC family member *Zic1* was not induced upon vanillin treatment (Figure S3i).These results strongly recommend ZIC3 as an inducer of TH and DAergic identity to cells in OB and could possibly play a key role in olfactory learning and discrimination.

### ZIC3 is essential for generation of TH expression in OB like neurons derived from mESCs

In order to test the requirement of *Zic3* for TH expression in an independent model system, we selected mESC based *in vitro* differentiation. mESCs were initially converted to embryoid bodies and further allowed to acquire neural patterning. After providing OB related cues (Figure 3a) the cells were characterized for expression for OB specific markers. A gradual increase in OB markers *Th, Gad65, Er81* and *Nurr1* was observed. *Map2*-a terminally differentiated neuron marker was also up-regulated along with a concomitant down-regulation of pluripotency marker *Oct4* (Figure 3b). These cells also co-expressed TH with GAD65, MAP2, TUJ1 (Figure S4a) and stained positive for OB enriched transcription factors ER81 and NGF1β (Figure S4b). This proved the establishment of an authentic differentiation system to test the role of ZIC3. Upon inhibition of *Zic3* using a specific shRNA, we could see an evident down-regulation of *Th* expression, but not the expression of dopamine cassette genes *Aadc, Vmat2* and *Gch1* (Figure 3c). Besides these, other OB markers *Er81* and *Ngf1b* also remained unchanged. Further protein analysis using western blot (Figure 3d.i), immunofluorescence (Figure 3d.ii) and flow cytometry (Figure 3d.iii) also revealed reduction in TH expression (Figure 3d.iv) upon *Zic3* knockdown. Thus, using two independent cellular models, our study advocates a novel role for ZIC3 in governing the expression of TH in OB DA neurons.

**Figure 3:**
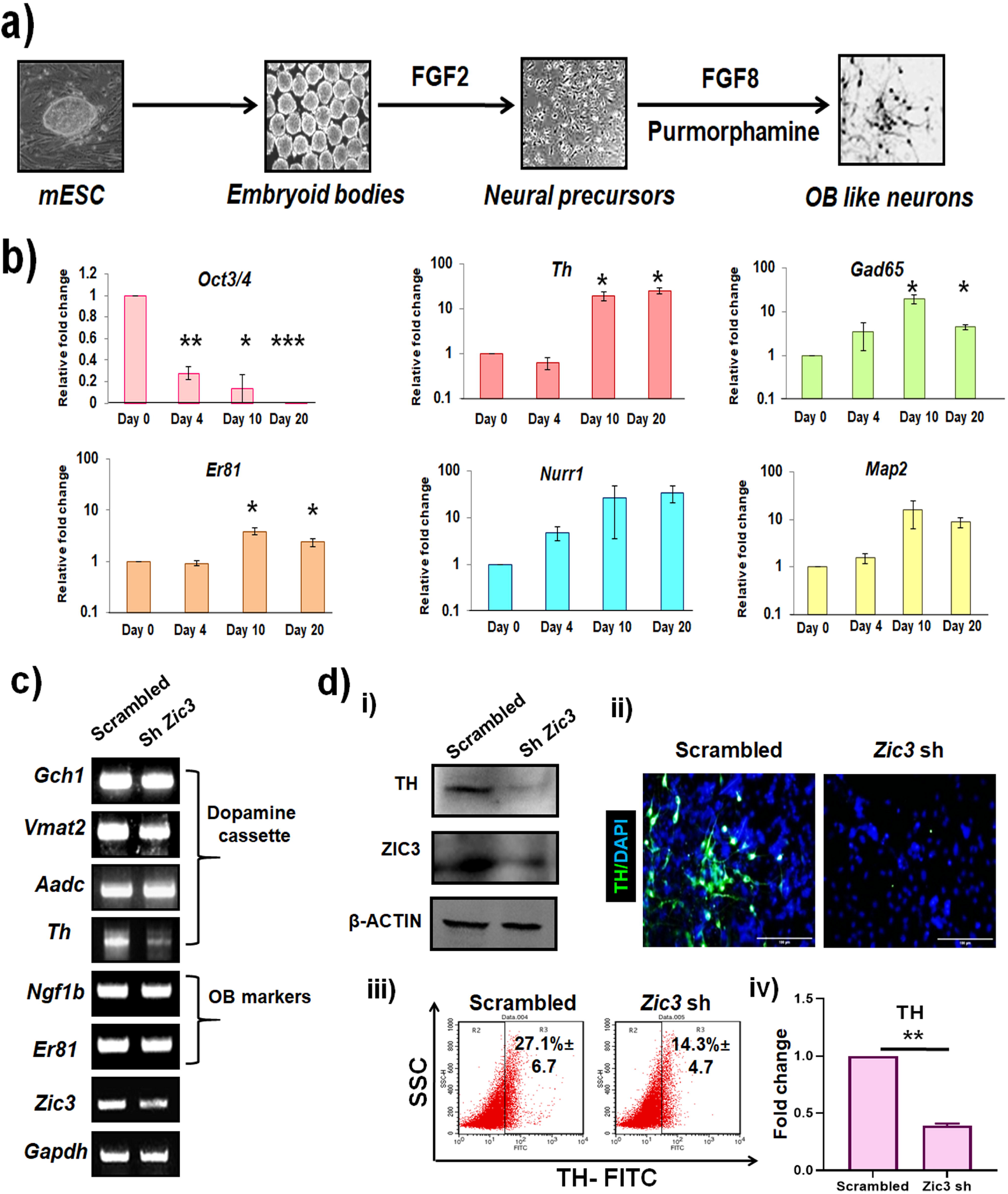
ZIC3 is essential for generation of OB DA-like neurons from mESC. (a) Schematic representation of mESC to OB neuron differentiation, b) Transcript analysis of pluripotent marker *Oct4* and various OB neuronal markers during different days of differentiation, c) mRNA analysis showing specific reduction in *Th* expression upon *Zic3* knockdown in OB differentiation of mESCs, (d) western blot (i), immunofluorescence (ii) and flow cytometry (iii) analysis of TH expression and its quantification (iv) in OB like neurons derived from mESCs in presence and absence of *Zic3*. Scale bar represents 100 µm. Mean+/-SE of biological triplicates, * p≤ 0.05, ** p ≤ 0.01, *** p ≤ 0.001.

### ZIC3 and ER81 co-regulate TH expression in OB DA neurons

After establishing a positive correlation between ZIC3 and TH in OB DA neuron generation we intended to understand the underlying mechanism. Initially, we performed protein-DNA interaction studies to determine if ZIC3 directly binds to *Th* promoter. Though the bio informatics analysis did not show any ZIC3 consensus binding site on *Th* promoter, we went ahead to perform chromatin immunoprecipitation (ChIP) assay to verify the results. Upon performing targeted PCR with the immunoprecipitated DNA for different regions of *Th* promoter with overlapping primers, surprisingly, we identified the occupancy of ZIC3 encompassing (+7 to -172 bp) region on *Th* promoter using both qPCR (Figure 4a.i) and semi-quantitative PCR (Figure 4a.ii). After sequencing the stretch of the promoter region occupied by ZIC3, we found the encompassed region to correlate with the binding site of protein ER81 (Figure 4b.i)-previously implicated in positive regulation of *Th* expression in mouse OB (Cave et.al., 2009). We performed genetic knockdown of ER81 (Figure S5 a.i) and observed significant reduction in *Th* mRNA (Figure S5 a.ii) and protein (Figure S5 b), thus supporting the previous study of involvement of ER81 in TH regulation (Cave et.al., 2009). The occupancy of ER81 was re-confirmed by performing ER81 ChIP, which showed an enrichment in the same region as that of occupancy of ZIC3 (Figure 4b.ii). This led us to hypothesize a probable physical interaction between ZIC3 and ER81 on *Th* promoter. To answer this, we cloned ZIC3 and ER81 in highly immunoreactive tags GST and FLAG respectively. Co-immunoprecipitation assay revealed a direct interaction between GST and FLAG (Figure 4.c.i). Alternatively, immunoprecipitation (forward and reverse) was performed using mouse OB tissue protein and determined a direct interaction between ZIC3 and ER81 (Figure 4.c.ii). This interaction was true only for ZIC3 but not ZIC1 (Figure 4.c.iii) thus suggesting the specificity of the phenomenon. Functional analysis using *Th*-luciferase assay showed a gradual increase in its expression upon transfection of HEK293T cells with increasing concentration of *Zic3* plasmid thus ascertaining direct regulation of *Th* by ZIC3 (Figure 4d.i). Extending the observation to OB primary neurons, we found inhibition of *Zic3* reduced the *Th*-luciferase by 30%, which was rescued by *Zic3* OE (Figure 4d.ii). Taking into consideration the interaction between ZIC3 and ER81 in *Th* regulation, we over-expressed ZIC3, ER81 individually and in combination and observed highest activation of *Th*-promoter when both the TFs were combined (Figure 4d.iii). This result conveyed the ZIC3 mediated TH regulation to be via its interaction with ER81.

**Figure 4:**
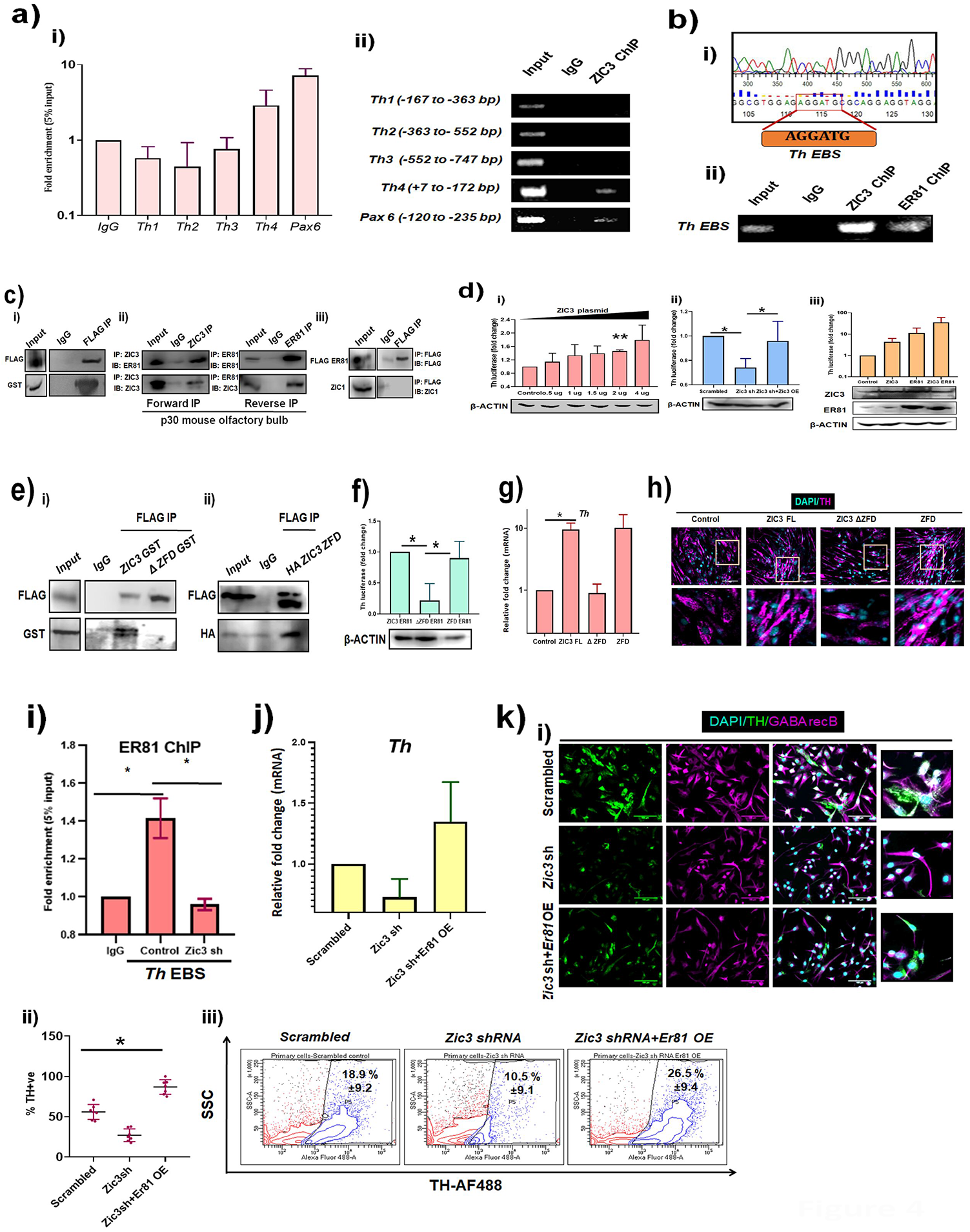
ZIC3 and ER81 interact with each other to regulate TH expression in mouse OB. (a) ChIP analysis showing ZIC3 occupancy on various regions of mouse *Th* promoter using region specific primers by (i) qPCR and (ii) conventional PCR. *Pax6* was used as a positive control for ZIC3 binding, b) Sanger sequencing result (i) and ChIP validation (ii) showing binding of ZIC3 and ER81 to *Th* promoter region in mouse OB, c) Co-immunoprecipitation analysis in HEK-293 cells overexpressing immunoreactive tags FLAG -ER81 and ZIC3-GST. Immunoprecipitation was performed using FLAG antibody and immunoblot performed using GST antibody. ii) Endogenous protein interaction between ZIC3 and ER81 was confirmed in mouse OB by forward immunoprecipitation using ZIC3 primary antibody for immunoprecipitation and immunoblot using primary ER81 antibody and by reverse immunoprecipitation using ER81 primary antibody for immunoprecipitation and immunoblot using ZIC3 primary antibody, iii) co-immunoprecipitation showing lack of interaction between ZIC1 and FLAG-ER81, d) i) Analysis of activation of *Th* promoter-luciferase in presence of increasing concentration of *Zic3* plasmid, β-actin used as loading control, ii) *Th* – promoter analysis in *Zic3* shRNA and simultaneous over-expression conditions, β-actin used as loading control and iii) in individual over-expression of ZIC3 and ER81 and in combined overexpression of ZIC3 and ER81 conditions. Western blots depicts the overexpression of ZIC3 and ER81 in their respective conditions and β-actin used as loading control. (e) Western blots showing interaction between ZIC3-FL and ER81 but not the ZFD deletion mutant (ΔZFD) (i) and overexpression of Zinc finger domain (ZFD) showed it to be sufficient to interact with ER81 (ii), (f) Analysis of activation of *Th* promoter-lucifearse upon ectopic expression of ZIC3-FL, ΔZFD and ZFD along with FLAG-ER81, β-actin used as loading control. (g) Transcript and protein (h) expression of TH in OB DA primary neurons over-expressing ZIC3-FL, ΔZFD and ZFD plasmids in ΔZFD condition, (i) ChIP analysis using ER81 antibody in OB DA neurons harboring scrambled and *Zic3* shRNA constructs, (j) Relative mRNA levels of *Th* in OB DA neurons with *Zic3* shRNA and *Er81* over-expression constructs, (k. i) OB DA primary neurons with *Er81* over-expression combined with *Zic3* inhibition conditions showing modulation in TH expression using immunofluorescence and its quantification (k. ii), and flow cytometric evaluation of TH expression (k. iii). Scale bar represents 100 µm. Mean+/-SE of biological triplicates, * p≤ 0.05, ** p ≤ 0.01, *** p ≤ 0.001.

Despite appreciable data on involvement of ZIC3 in various developmental processes its interactome is poorly built and the functions carried out by different domains of *Zic3* gene remain elusive. The present study revealed ER81 as a novel interacting protein partner of ZIC3 in OB DA neuron development leading to further questions on *Zic3* domain involved in this interaction. ZIC3 consists of five highly conserved zinc finger domains (ZFD) and studies in the past have advocated the involvement of ZFD in nuclear localization (Ali et.al. 2012), regulation of ZIC3 protein stability (Chhin et.al. 2007) and protein-protein interaction (Pourebrahim et al., 2001, Mizugishi et al., 2011). To determine the involvement of ZFD in the context of DA fate commitment, we generated deletion mutants of ZIC3 lacking one or more zinc fingers, in a GST backbone. The immunofluorescence images showed while the mutants ΔZF5, ΔZF4-5, ΔZF3-5 were able to localize to the nucleus, ΔZF2-5 and ΔZFD accumulated in the cytoplasm due to loss of nuclear localization signal (NLS) (Figure S6 a). Interaction studies showed that only ZIC3-FL but neither the complete deletion of ZFD nor the nuclear localized partial ZFD mutants were able to interact with ER81 (Figure 4 e.i and S6 b). Interestingly, just the over-expression of ZFD was sufficient to physically associate with ER81 as shown by co-immunoprecipitation (Figure 4e. ii). Extrapolating the results of ZFD interaction studies, we identified that ZIC3 lacking ZFD failed to drive the *Th*-luciferase due to lack of its interaction with ER81. However, a mere substitution with ZFD region along with FLAG ER81 was sufficient to prompt *Th*-luciferase expression to the extent of the expression up-regulated by ZIC3 FL and ER81 together (Figure 4f). To understand whether the role of interaction domains hold true in primary neurons, we overexpressed ZIC3 -FL, ΔZFD and ZFD and observed that only full length ZIC3-GST clone was able to up-regulate *Th* expression by about 10 folds during OB DA neuron differentiation but ΔZFD failed to drive the *Th* expression (Figure 4g). Reconstitution of just the ZFD could also lead to a 10 fold increase in *Th* mRNA similar to the FL clone; albeit the over-expression of ZIC3 full length or ZFD was sufficient to rescue the TH protein production in these neurons whereas the ΔZFD failed to have an effect on TH expression (Figure 4h).

Though our results so far advocated a role for ZIC3 and ER81 association in OB DA neurons, the indispensability of ZIC3 for ER81 mediated regulation of *Th* remained unanswered. To address this, a ChIP analysis was performed and observed that upon inhibition of *Zic3* in OB DA primary neurons, binding of ER81 to *Th* promoter was considerably decreased (Figure 4i), thus projecting ZIC3 as a co-regulator which dictates the binding of ER81 to *Th*-promoter. Also, in absence of ZIC3, over-expression of ER81 rescued the expression of *Th* transcript and protein to the level of that in scrambled control (Figure 4j and 4k.i, ii). Flow cytometric analysis also showed enhanced TH +ve cells in ER81 over-expression condition, despite *Zic3* knockdown (Figure 4k. iii). On the other hand, supplementing *Zic3* in cells lacking *Er81* could not elicit a rescue of *Th* expression (Figure S7 a). Thus, all these data together established a novel ZIC3-ER81-*Th* axis in DA fate specification.

### ZIC3 regulates TH expression in MB DA neurons

Besides OB, DA neurons are present in different regions of brain including Substantia nigra pars compacta (SNpc) and Ventral tegmental area (VTA) of midbrain. Our results so far had strongly advocated crucial role for ZIC3 in regulating the DA identity of OB interneurons by influencing TH expression. To test if the same holds true for MB DA neurons we initially sorted TH positive neurons from 4-week-old mouse MB and stained for ZIC3. Approximately 27% of the cells co-expressed TH and ZIC3 (Figure 5a). To test the requirement of ZIC3 in MB DA neurons, primary neurons from E 12.0-E 14.0 mouse MB were derived by generating neurospheres and further differentiating them to DA like fate. While the neural progenitors were positive for SOX1 and NESTIN, their differentiated counterparts expressed TUJ1 and TH (Figure S8 a) thus authenticating DA like signatures. Upon *Zic3* inhibition using a shRNA, there was a significant reduction in *Zic3* expression (Figure S8 b) with a concomitant 80% reduction of *Th* mRNA with respect to scrambled control (Figure 5b). Immunofluorescence staining (Figure 5c. i and ii) and flow cytometry analysis also showed about 16% down-regulation of TH protein levels in *Zic3* knockdown condition with respect to the scrambled control (Figure 5d. i and ii).

**Figure 5:**
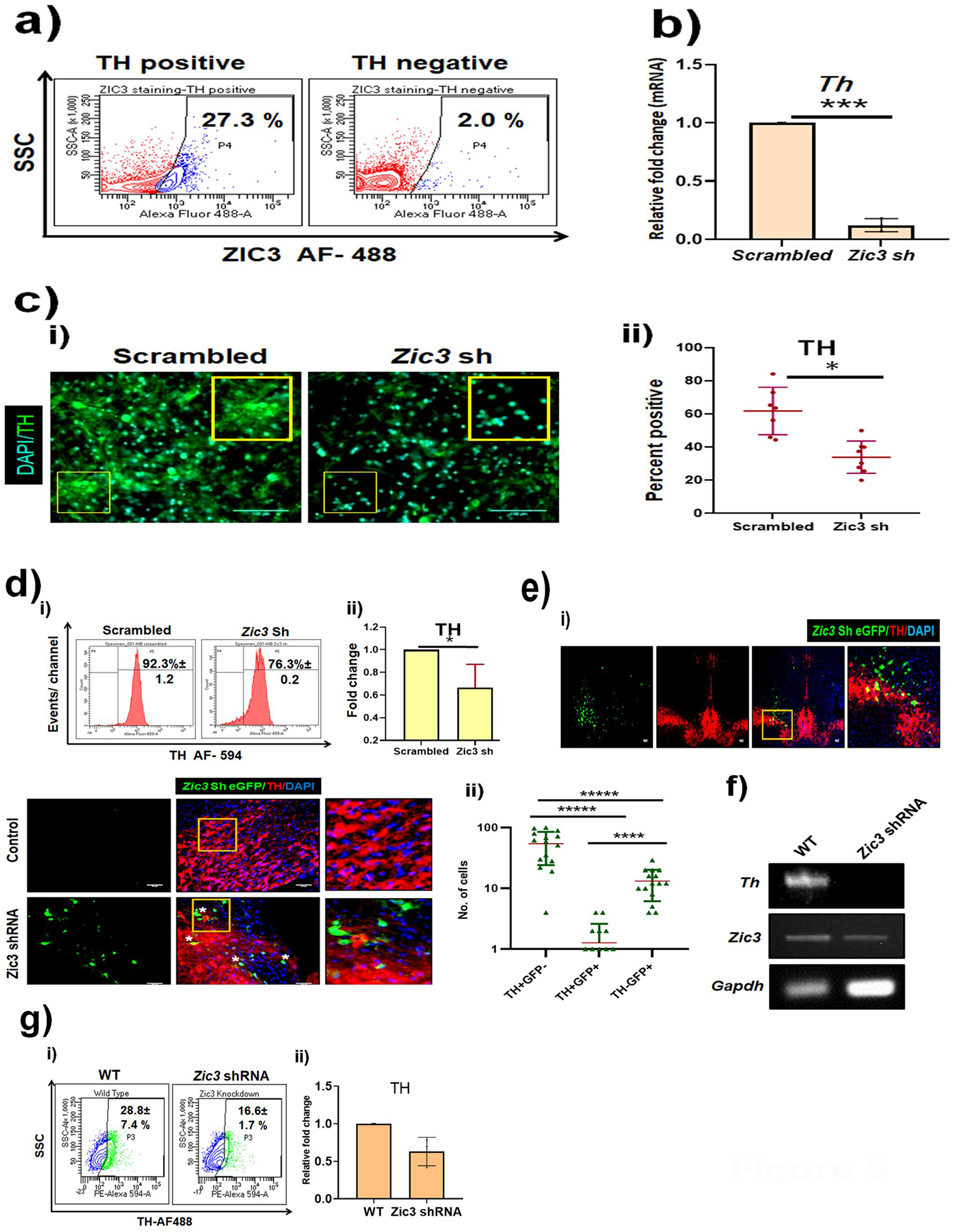
ZIC3 positively regulates TH expression in MB DA neurons. (a) Flow cytometric analysis of ZIC3 expression in TH positive and negative cells sorted from mouse MB, (b)Transcript and (c. i) protein analysis of Th, followed by quantification (c. ii) of primary MB DA neurons cultured in presence or absence of Zic3 shRNA, (d. i) flow cytometry analysis and its quantification (d. ii) upon Zic3 knockdown in MB DA primary neurons, e. i) *in vivo* expression analysis of TH in mouse MB upon stereotactic injection of Zic3 shRNA-eGFP and (ii) quantification of TH+/GFP-, TH+/GFP+ and TH-/+GFP cells. (f) transcript analysis and flow cytometry (g. i) for TH in *Zic3* knockdown mouse MB cells followed by its quantification (g. ii). Scale bar represents 100 µm. Mean+/-SE of biological triplicates, * p≤ 0.05, *** p≤ 0.001 and ***** p ≤ 0.0001.

In order to comprehend this regulation in an *in vivo* scenario, we performed stereotactic delivery of *Zic3* shRNA tagged to eGFP in 7 months old mice MB region. Post ten days of injection, the animals were sacrificed, and the MB region was stained for the expression of GFP and TH (Figure 5e. i). While 56%±7.8 cells lacking *Zic3* shRNA were TH+ve in both SNpc and VTA regions, only 1.2%±0.3 of those with *Zic3* shRNA were GFP+ve and TH-ve (Figure 5e. ii). Further, transcript analysis showed a drastic reduction in *Th* expression in the midbrain region stereotactically injected with *Zic3* shRNA (Figure 5f). In order to quantify the reduction in TH positivity, we performed flow cytometric staining using midbrain cells from control and *Zic3* shRNA injected mice and found an approximate 12% reduction in the TH positive cells upon *Zic3* knockdown (Figure 5g i and g. ii). This result demonstrated ZIC3 to act as a positive regulator of TH in mouse MB, similar to our observations in OB DA pool.

### ZIC3 switches molecular partners in OB and MB DA neurons to regulate TH expression

Having discovered an important role for ZIC3 in regulating *Th* expression and consequent DAergic fate of both OB and MB like neurons, we were intrigued to know whether ZIC3 adapts a similar mechanism as observed in OB. Corroborating the previous observations (Wang and Turner, 2010), flow cytometric analysis failed to detect the expression of ER81 expression in MB (Figure 6a). ChIP analysis in MB tissue lysate showed lack of ZIC3 binding to *Th* promoter region harboring ER81 binding site (Figure 6b). This suggested a possible alternate mechanism that could be adopted by ZIC3 to regulate TH in MB to compensate for the absence of its molecular partner ER81. In order to delineate the alternate route, we performed an *in silico* promoter analysis to find ZIC3 consensus binding site on various MB specific TFs namely *Otx2, Lmx1a/b, Engrailed, Foxa2, Nurr1* and *Pitx3*. Amongst these DA regulators, only *Pitx3* promoter harbored a ZIC3 binding site (Figure 6c). To determine if ZIC3 indeed bound to *Pitx3* promoter, we carried out a ChIP assay in 4 weeks old mouse MB lysate and found approximately 2-fold enrichment in ZIC3 binding to *Pitx3* promoter (Figure 6d.i). Interestingly, this binding was restricted to MB and was undetectable in OB tissue lysate (Figure 6d.ii). ChIP analysis in OB and MB lysates showed ZIC3 to preferentially bind *Th* EBS in OB but not in MB while it chooses to occupy *Pitx3* promoter in

**Figure 6:**
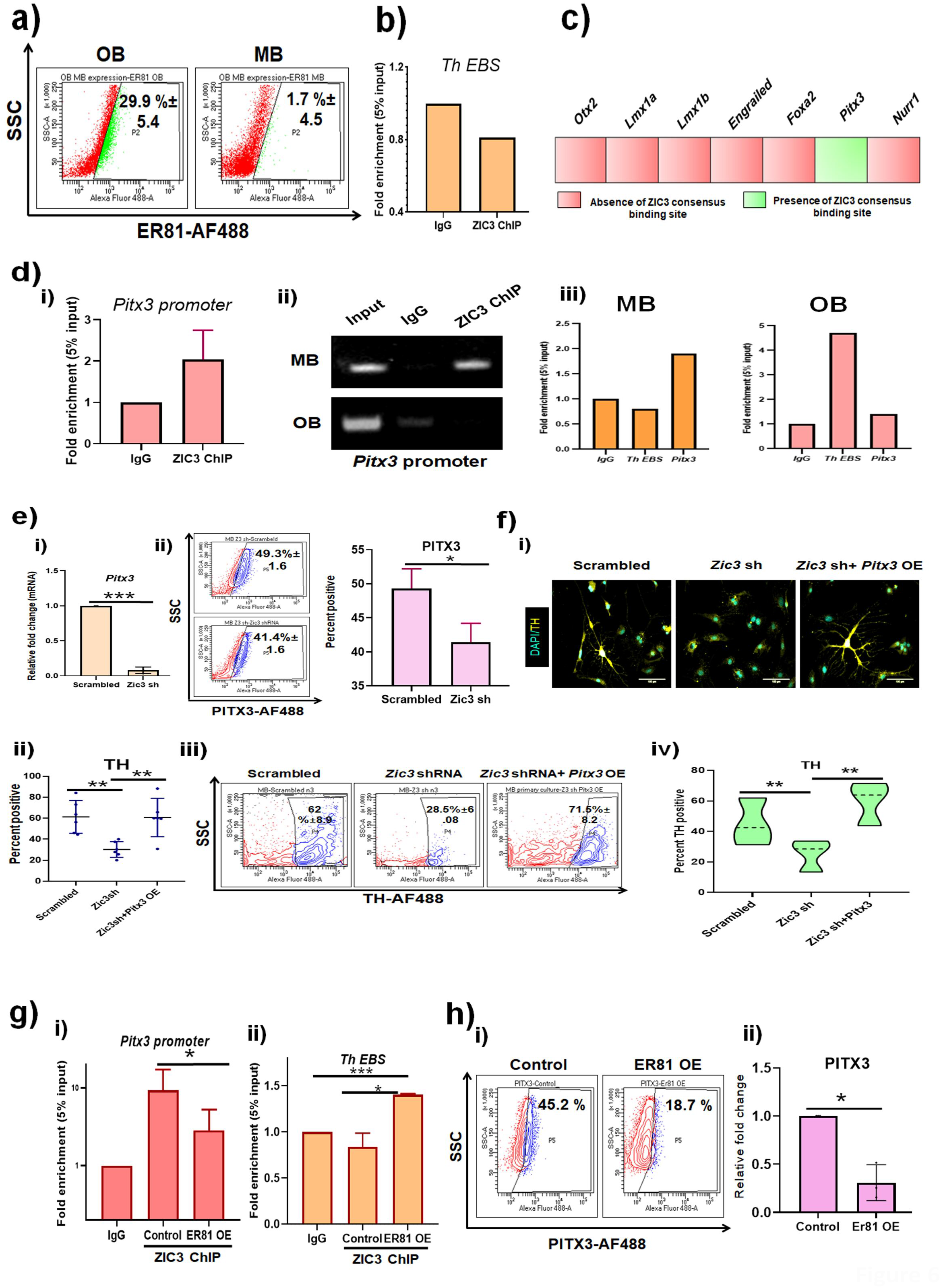
ZIC3 differentially regulates TH in mouse OB and MB by switching between ER81 and Pitx3. (a) Flow cytometric analysis of ER81 expression in mouse OB and MB, (b) ChIP analysis showing absence of ZIC3 binding on Th promoter region where ER81 binds, (c) Schematic representation of key midbrain specific gene promoters with presence and absence of ZIC3 consensus binding site in the region of -1000 bp from the transcript start site, (d) ChIP analysis showing occupancy of ZIC3 on *Pitx3* promoter in mouse OB tissue lysate as analysed by conventional PCR (d. i) and qPCR (d. ii). (d. iii) analysis of ChIP using ZIC3 primary antibody showing ZIC3 binding to *Pitx3* promoter specifically in mouse MB but not OB. Reduction in *Pitx3* mRNA (e. i) and *Pitx3* protein expression (e. ii) upon *Zic3* inhibition in MB primary neurons, (f) immunofluorescence for TH (f. i) and its quantification (f. ii), and flow cytometric analysis (f. iii) and its quantification (f. iv) of TH positive cells in MB DA neurons upon *Zic3* inhibition along with *Pitx3* over-expression. (g) ChIP analysis using ZIC3 antibody showing reduced binding of ZIC3 to *Pitx3* (g. i) and concomitant increase in occupancy of ZIC3 on *Th* (g. ii) promoter in primary MB neurons. h) Flow cytometric analysis (i) of PITX3 expression and its quantification (ii) in MB primary neurons over-expressing ER81. Scale bar represents 100 µm. Mean+/-SE of biological triplicates, * p< 0.05, ** p < 0.01, *** p< 0.001.

MB with an absence of any such association in OB (Figure 6d. iii). Also, inhibition of *Zic3* in MB primary neurons led to significant down-regulation of *Pitx3* mRNA (Figure 6e.i) and protein (Figure 6e.ii). While knockdown of *Zic3* abrogates *Th* expression in primary MB DA neurons, over-expression of *Pitx3* along with *Zic3* shRNA retains TH expression as shown by immunofluorescence (Figure 6f. i and ii). Flow cytometry also reiterated *Pitx3* as the downstream effector of ZIC3 in TH regulation owing to the retention of TH expression in cells subjected to *Zic3* loss of function (Figure 6f.iii and 6f.iv). This result conclusively established *Pitx3* as the down-stream target of ZIC3 in MB and as a novel interaction operating in DA fate specification.

Our results so far reveal a choice for ZIC3 in selecting different interacting partners in OB and MB to orchestrate *Th* regulation. This intrigued us to examine the molecular interaction pattern upon external supplementation of ER81 in MB cells that inherently lacked it. Forced over-expression of ER81 in MB primary neurons led to decreased binding of ZIC3 to *Pitx3* promoter (Figure 6g.i) with a collateral establishment of its association with *Th* promoter similar to the scenario in OB DA neurons (Figure 6g.ii). This also translates to modulation in PITX3 expression levels wherein, only 18.7% of the cells were PITX3 positive upon *Er81* over-expression as against 45.2% positive cells in control differentiation (Figure 6h.i) which accounted for about 60% down-regulation in PITX3 expression (Figure 6h.ii). All these results advocate a switch in molecular partners of ZIC3 to ensure spatially different *Th* regulation (Figure 7).

**Figure 7:**
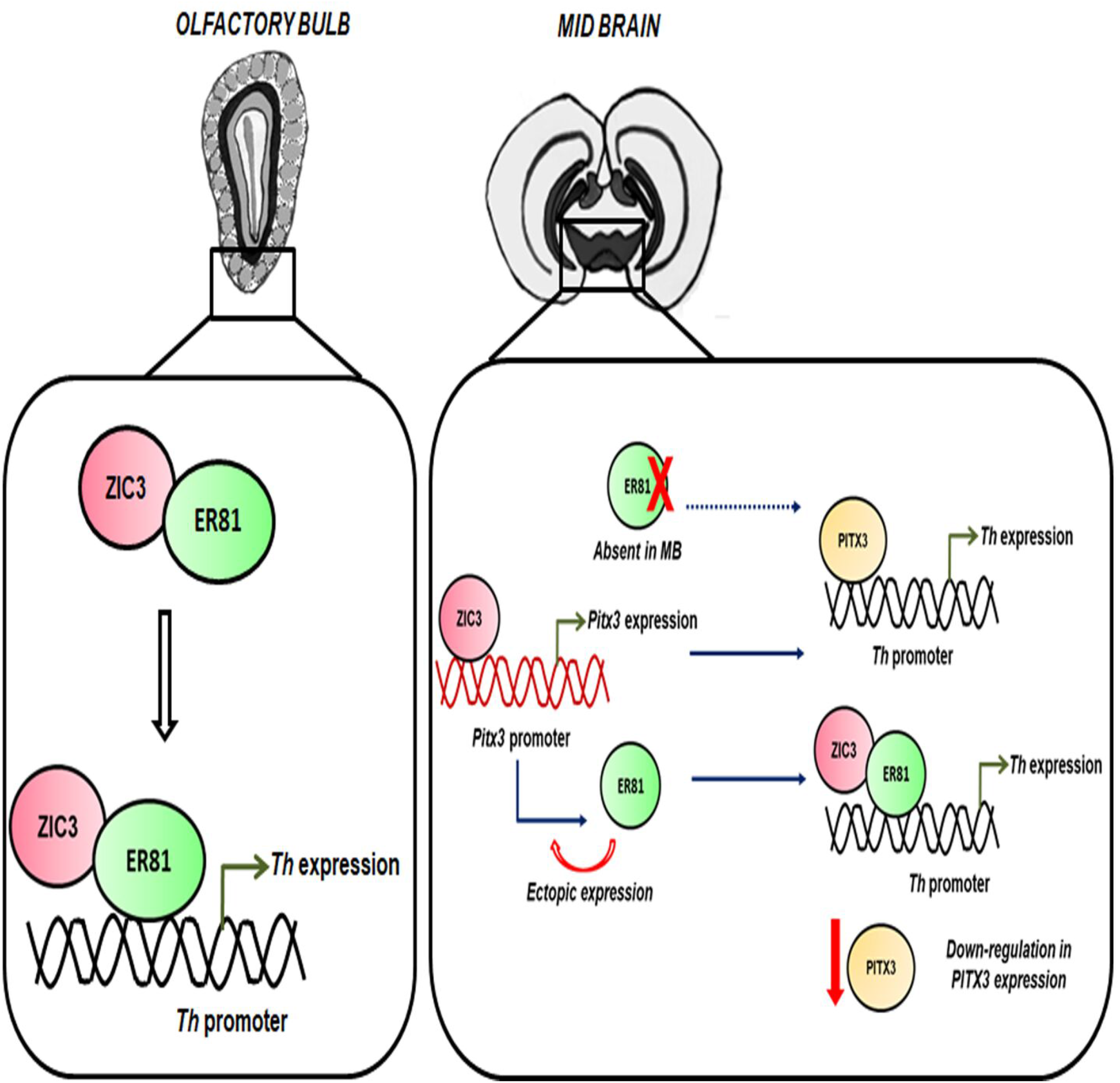
Schematic representation of the bimodal regulation of TH by ZIC3 in mouse OB and MB. In the olfactory bulb, ZIC3 collaborates with ER81 and co-regulates *Th* expression by binding the promoter. While in the midbrain, due to absence of ER81, ZIC3 binds to Pitx3 promoter and drives the expression. PITX3 in turn leads to the expression of *Th*. Upon ectopic expression of ER81 in MB DA primary neurons, ZIC3 switches the molecular partner and binds along with ER81 to *Th* promoter, which further leads to down-regulation of PITX3 expression.

## Discussion

In this study we present evidence of a protein ZIC3 as a novel intrinsic determinant of DAergic neuronal fate in OB and MB. Targeted manipulations of ZIC3 expression in the OB and MB primary neurons showed the importance of ZIC3 in the generation of DAergic neurons. The observation was ascertained by using primary neurons, ESC derived OB neurons and *in vivo* mouse model system. Notably, this role is specific to ZIC3 among ZIC family members, as the knockdown of ZIC1 failed to modulate the expression of *Th*. Attempt to understand the mechanism demonstrated ZIC3 to utilize different modes of regulation in OB and MB to dictate the DA fate. In OB, ZIC3 forms higher order protein complexes and functionally interacts with ER81, a previously known regulator of DAergic specification in OB. On the contrary, in absence of ER81 in MB derived primary neurons, ZIC3 directly occupies *Pitx3* promoter and indirectly regulates the expression of *Th*. Together, these findings assign a novel function to TF ZIC3 and describe an interesting region specific biphasic mechanistic model to steer DAergic cell fate decision.

Transcriptional codes directing midbrain (Ang S.L., 2006) and forebrain DAergic identities are categorical (Allen Z.J., et.al 2007). More often, the neurochemical and morphological properties of these neurons are governed by the spatial source of stem cells from which they originate (Angelova A., et.al 2018). OB interneurons are majorly contributed from the lateral ganglionic eminence (LGE) (Stenman J., et.al 2003) and septum (Qin S., et.al 2017), from embryonic day 12 (E12) until birth and in adult it is arriving from SVZ (Lois C., et.al 1996). By the virtue of their place of birth and migratory path, these cells acquire various molecular codes that facilitate DA fate establishment (Hack M.A., et.al 2005; Diaz-Guerra., et.al 2013). These molecular codes, especially TFs, by region specific variability in their expression level impart specific cell fate. While studies in the past have reported ER81 expression in OB DA interneurons globally (Saino-Saito S., et.al 2007), ZIC1 and ZIC2 expressions were shown to be exclusive of TH positive cells and restricted to CalR positive GABAergic interneurons. Also, it was shown that ZIC1and ZIC2contribute to the generation of CalR-positive neurons by suppressing DAergic fate (Tiveron M.C., et.al 2017). Recently, Quin et al., showed using ZIC3-LacZ model system that *Zic3* expressing cells originate from the septum and migrate towards the OB to populate several cell types including the DA interneurons (Qin S., et.al 2017). To convincingly show that the regulation of *Th* by ZIC3 is a cell autonomous phenomenon, we performed spatial distribution studies, which suggested a sub-population of TH positive neurons, to express ZIC3-accounting for about one third of the total DA neuron population. This sub-set of neurons plausibly suggests the septal origin of these cells. It would be interesting to further study the expression pattern of ZIC3 in SVZ, RMS and its requirement for maintenance of NSC pool, neuroblasts migration and by and large, adult neurogenesis.

Our experiments reveal that ZIC3 has a positive correlation with TH expression, and it leads to compromised DAergic identity upon inhibition. Our results corroborate well with the previously reported retarded OB development in compound mutants of ZIC1 and ZIC3 (Inoue T., et.al 2007). DAergic interneurons of OB are distinct from those of other regional identities due to their partial GABAergic nature (Gall C.M., et.al 1987). While ZIC3 modulates the DAergic identity of the cells, the GABAergic phenotype remains unaltered. Thus, ZIC3 might be a cue to specifically drive TH expression and cajole them towards DAergic lineage. Additionally, the current study advocates the ability of ZIC3 to override the DAergic cues-FGF2, FGF8 and SHH in facilitating TH expression. Contribution of ZIC3 to DA fate strongly hints towards its plausible involvement in their lineage determination. As the major source of these DAergic neurons of OB is adult neurogenesis, a role for ZIC3 in this event cannot be ruled out. Ablation of ZIC3 expression post differentiation also led to down-regulation of TH expression, strongly indication the role of ZIC3 in maintaining the identity of TH+ve neurons apart from their generation. As we did not notice any significant cell death upon ZIC3 abrogation, the compromise in TH expression upon ZIC3 inhibition probably did not lead elimination of these neurons and perhaps led to an alternative neurochemical identity, which is yet to be deciphered.

Previous studies provided insights into the implications of ZIC3 on neural progenitor expansion (Inoue T., et.al 2007) and fate commitment. ZIC3 is usually viewed as a DNA distal binding element and also acts as a co-activator for other TFs (Winata C., et.al 2013; Mizugishi K., et.al 2001). While ZIC3 binds to *Nanog* promoter region to promote stemness in embryonic stem cells, it collaborates with TFs such as SOX2 (Lim L.S., et.al 2010) in the context of pluripotency and ESRRB to synergistically switch the metabolic fates during cellular reprogramming (Sone M., et.al 2017). Mechanistic investigation into ZIC3 mediated regulation of OB DA neuronal fate showed an absence of consensus binding site for ZIC3 on *Th* promoter. However, a DNA binding enrichment was seen in a region of *Th* encompassing a DNA sequence devoid of previously reported ZIC3 recognition site. Sanger sequence demonstrated the presence of ER81 binding site in the above said stretch of DNA thus suggesting an interaction between ZIC3 and ER81. Interaction studies revealed a direct association between ZIC3 and ER81 in OB tissue thus providing satisfactory explanation to ZIC3 mediated regulation of TH. We also demonstrate zinc finger domains of ZIC3 to be sufficient for interaction with ER81 and subsequent regulation of *Th* expression. The functional significance of the interaction between ZIC3 and ER81 is appreciated by the reduced occupancy of ER81 on the *Th* promoter in the absence of ZIC3. Surprisingly, the compromised expression of TH in absence of ZIC3 was rescued by over-expression of ER81, but not conversely when ZIC3 was over-expressed in absence of ER81 (figure S7 a). In-depth understanding showed ER81 enhance the expression of *Zic3* even under the influence of *Zic3* shRNA (Figure S7 b) thus enabling retention of *Th* expression. ChIP analysis showed the direct binding of ER81 on *Zic3* promoter (Figure S7. c) thus satisfactorily explaining the ER81 mediated rescue of TH upon *Zic3* inhibition. In all, by utilizing interaction and functional assays, we reveal a novel association between ZIC3 and ER81 in OB DA neurons.

Few TFs in a spatially restricted manner regulate the DAergic neurogenesis (Remesal L., et.al 2020; Backman C., et.al 1999; Cave J.W., et.al 2010; Hwang D.Y., et.al 2003) and it is imperative to comprehend these events as they could provide valuable answers to differential disease response of DA neurons. While PITX3 is actively engaged in regulating TH expression in midbrain (Smit S.M., et.al 2006), its expression in OB is undetectable. NURR1, a TF which is essential for maintenance and differentiation of ventral mesencephalic DAergic precursor cells to mature DA neurons (Kadkhodaei B., et.al 2009; Sakurada K., et.al 1999), it is present in OB but lacks the ability to regulate the TH expression (Liu N., 1999). Though ER81 is expressed in the cortex during the embryonic day E15 -P14 and involved in the differentiation of the layer 5 cells of the cortex, no detectable expression is observed in the adult cortex (Yoneshima H., et.al 2006). ER81 is also not expressed in the ventral part of the midbrain in both embryonic and adult life (Wang and Turner 2010). While the present study showed ZIC3 to interact with ER81 and finetune the ER81 mediated TH expression in OB, the lack of ER81 and its incapability to influence dopamine pathway (Wang and Turner 2010) in MB compelled us to introspect the alternative mechanism of ZIC3 mediated TH regulation in MB. *In silico* screening for ZIC3 binding to different promoters of TFs involved in TH regulation accentuated PITX3 as a prominent candidate gene. Further, ChIP assay confirmed *Pitx3* to be the downstream target of ZIC3. Reports have established the importance of tight transcriptional regulation of PITX3 in DA identity and their survival. Our results bring into fore, ZIC3 as a novel upstream regulator of PITX3. Having established ZIC3 mediated regulation of TH in OB and MB DA neurons, we set out to determine its preference to molecular partners. An evident displacement of ZIC3 from *Pitx3* promoter following its association with ectopically expressed ER81 in MB probably hints towards higher affinity of ZIC3 to ER81.

Highly complicated and distinct molecular programs that operate in DA neurons might obligate ZIC3 to control DA fate, region specifically. Thus, though ZIC3 lacks the ability to directly bind to *Th* promoter and drive its expression, through its molecular partners and mechanisms ZIC3 enables the fine tuning of *Th* expression.

Early TFs of OB lineage development, such as PAX6 are involved in DA neuron survival (Agoston Z., et.al 2014) whereas NEUROD1 is a key for the terminal maturation of these neurons (Boutin C., et.al 2010). ETS family member ER81 regulates DA fate commitment by directly driving *Th* expression (Flames N., et.al 2009; Cave J.W., et.al 2010). Recently, a TF PBX1 is shown to assist in the end point selection of OB DA neurons (Remesal L., et.al 2020). However, none of these TFs have been shown to differentially regulate the TH +ve neuron pool outside of OB. Interestingly, *Zic3* as reported in the current study, regulates DA neurons in spatially defined micro-domains by the virtue of its molecular partner switching mechanism. This novel bimodal control of TH expression could contribute however minimally, to render these neurons distinct from one another.

It would also be intriguing to know how this role of ZIC3 can be extrapolated to several other pockets of DA neurons throughout the brain. Studying these events could be clinically relevant as OB and MB DA neurons respond differently to neurodegenerative insults like those in PD. In addition to regulating TH during the course of differentiation, ZIC3 also enhances TH expression upon olfactory stimulation. In retrospect, ZIC3 expression is also enhanced in the OB of mice housed in odor enriched environment in comparison to those under limited exposure to odor stimulants. This interesting observation suggests that ZIC3 could probably be a marker to assess the state of olfactory perception. As hyposmia is one of the earliest symptoms in PD patients (Xiao Q., et.al 2014), any contribution of ZIC3 to olfactory perception could be instrumental in ascertaining its importance in DA neuron function and disease predisposition. Our study reveals a previously unknown molecular program involving ZIC3 and ER81 during OB DA development. This information would facilitate engineering pluripotent stem cell derived and transdifferentiation based approaches to derive DAergic neurons for probable cellular therapies to mitigate DA related disorders.

## ACKNOWLEDGEMENT

This study was supported by grants from Department of Biotechnology, India (BT/PR 6165/GBD/27/369/2012, BT/PR8508/MED/30/1019/2013) to A.K. S.B. was supported by CSIR SRF (#09/1249 (0002) 2K19 EMRI). We acknowledge MAHE for providing intramural fund. We thank Professor Colin Jamora, IFOM-inStem Joint Research Laboratory, Institute for Stem Cell Science and Regenerative Medicine (inStem) for IHC related work, Dr. H. Krishnamurthy and the staff of Central Imaging and Flow Cytometry Facility (CIFF), NCBS, Bangalore for their help with FACS.

## AUTHOR CONTRIBUTIONS

Conceptualization, S.B, J.P., and A.K.; Methodology and Formal Analysis, S.B. and A.K.; Investigation, S.B., J.G., A. H., S.C.R.T and N.R; Writing – Original Draft, S.B., J.G., and A.K.; Writing – Review & Editing: S.B., J.P., S.C.R.T., N.R., and A.K.; Supervision, A.K.; Funding Acquisition, A.K.

## References

Agoston, Z., Heine, P., Brill, M.S., Grebbin, B.M., Hau, A.C., Kallenborn-Gerhardt, W., Schramm, J., Götz, M. and Schulte, D. 2014. Meis2 is a Pax6 co-factor in neurogenesis and dopaminergic periglomerular fate specification in the adult olfactory bulb. Development. 141:28–38.

Allen, Z.J., Waclaw, R.R., Colbert, M.C. and Campbell, K. 2007. Molecular identity of olfactory bulb interneurons: transcriptional codes of periglomerular neuron subtypes. J. Mol. Histol. 38: 517–525.

Ang, S.L. 2006. Transcriptional control of midbrain dopaminergic neuron development. 3499–3506.

Angelova, A., Tiveron, M.C., Cremer, H. and Beclin, C. 2018. Neuronal subtype generation during postnatal olfactory bulb neurogenesis. J. Exp. Neurosci. 12:1179069518755670.

Aruga, J., Nagai, T., Tokuyama, T., Hayashizaki, Y., Okazaki, Y., Chapman, V.M. and Mikoshiba, K. 1996. The Mouse Zic Gene Family: HOMOLOGUES OF THE DROSOPHILA PAIR-RULE GENE odd-paired (*). J. Biol. Chem. 271:1043–1047.

Aruga, J., Yokota, N., Hashimoto, M., Furuichi, T., Fukuda, M. and Mikoshiba, K. 1994. A novel zinc finger protein, zic, is involved in neurogenesis, especially in the cell lineage of cerebellar granule cells. J. Neurochem. 63:1880–1890.

Aruga, J., Yozu, A., Hayashizaki, Y., Okazaki, Y., Chapman, V.M. and Mikoshiba, K. 1996. Identification and characterization of Zic4, a new member of the mouse Zic gene family. Gene. 172: 291–294.

Bäckman, C., Perlmann, T., Wallén, Å., Hoffer, B.J. and Morales, M. 1999. A selective group of dopaminergic neurons express Nurr1 in the adult mouse brain. Brain. Res. 851:125–132.

Banerjee, A., Marbach, F., Anselmi, F., Koh, M.S., Davis, M.B., da Silva, P.G., Delevich, K., Oyibo, H.K., Gupta, P., Li, B. and Albeanu, D.F. 2015. An interglomerular circuit gates glomerular output and implements gain control in the mouse olfactory bulb. Neuron. 87:193–207.

Bonzano S, Bovetti S, Gendusa C, Peretto P, De Marchis S. Adult born olfactory bulb dopaminergic interneurons: molecular determinants and experience-dependent plasticity. 2016. Front. neurosci. 6;10:189.

Boutin, C., Hardt, O., de Chevigny, A., Coré, N., Goebbels, S., Seidenfaden, R., Bosio, A. and Cremer, H. 2010. NeuroD1 induces terminal neuronal differentiation in olfactory neurogenesis. Proc. Nat. Acad. Sci. 107:1201–1206.

Brill, M.S., Snapyan, M., Wohlfrom, H., Ninkovic, J., Jawerka, M., Mastick, G.S., Ashery-Padan, R., Saghatelyan, A., Berninger, B. and Götz, M. 2008. A dlx2-and pax6-dependent transcriptional code for periglomerular neuron specification in the adult olfactory bulb. J. Neuroci. 28:6439–6452.

Carrel, T., Herman, G.E., Moore, G.E. and Stanier, P. 2001. Lack of mutations in ZIC3 in three families with neural tube defects. Am. J. Med. Gen. 98:283–285.

Carrel, T., Purandare, S.M., Harrison, W., Elder, F., Fox, T., Casey, B. and Herman, G.E. 2000. The X-linked mouse mutation Bent tail is associated with a deletion of the Zic3 locus. Hum. Mol. Genet, 9:1937–1942.

Cave, J.W., Akiba, Y., Banerjee, K., Bhosle, S., Berlin, R. and Baker, H. 2010. Differential regulation of dopaminergic gene expression by Er81. J.Neurosci. 30:4717–4724.

Declercq, J., Sheshadri, P., Verfaillie, C.M. and Kumar, A. 2013. Zic3 enhances the generation of mouse induced pluripotent stem cells. Stem cells. Dev. 22:2017–2025.

Díaz-Guerra, E., Pignatelli, J., Nieto-Estévez, V. and Vicario-Abejón, C. 2013. Transcriptional regulation of olfactory bulb neurogenesis. Anat. Rec. 296:1364–1382.

Flames, N. and Hobert, O. 2009. Gene regulatory logic of dopamine neuron differentiation. Nature. 458:885–889.

Fritz, B., Kunz, J., Knudsen, G.P.S., Louwen, F., Kennerknecht, I., Eiben, B., Ørstavik, K.H., Friedrich, U. and Rehder, H. 2005. Situs ambiguus in a female fetus with balanced (X; 21) translocation–evidence for functional nullisomy of the ZIC3 gene?. Eu. J. Hum. Genet. 13:34–40.

Gall, C.M., Hendry, S.H., Seroogy, K.B., Jones, E.G. and Haycock, J.W. 1987. Evidence for coexistence of GABA and dopamine in neurons of the rat olfactory bulb. J. Comp. Neurol. 266:307–318.

Garnett, A.T., Square, T.A. and Medeiros, D.M. 2012. BMP, Wnt and FGF signals are integrated through evolutionarily conserved enhancers to achieve robust expression of Pax3 and Zic genes at the zebrafish neural plate border. Development. 139:4220–4231.

Gebbia, M., Ferrero, G.B., Pilia, G., Bassi, M.T., Aylsworth, A.S., Penman-Splitt, M., Bird, L.M., Bamforth, J.S., Burn, J., Schlessinger, D. and Nelson, D.L. 1997. X-linked situs abnormalities result from mutations in ZIC3. Nat. Genet. 17:305–308.

Hack, M.A., Saghatelyan, A., de Chevigny, A., Pfeifer, A., Ashery-Padan, R., Lledo, P.M. and Götz, M. 2005. Neuronal fate determinants of adult olfactory bulb neurogenesis. Nat. Neurosci. 8:865–872.

Hala, N., Ho, T., Johansson, O., Goldstein, M., Park, D. and Biberfeld, P. 1977. Transmitter histochemistry of the rat olfactory bulb. I. Immunohistochemical localization of monoamine synthesizing enzymes. Support for intrabulbar, periglomerular dopamine neurons. Brain Res. 126:455–474.

Halász, N., Johansson, O., Hökfelt, T., Ljungdahl, Å. and Goldstein, M. 1981. Immunohistochemical identification of two types of dopamine neuron in the rat olfactory bulb as seen by serial sectioning. J. Neurocyt. 10:251–259.

Havrda, M.C., Harris, B.T., Mantani, A., Ward, N.M., Paolella, B.R., Cuzon, V.C., Yeh, H.H. and Israel, M.A. 2008. Id2 is required for specification of dopaminergic neurons during adult olfactory neurogenesis. J. Neurosci, 28:14074–14087.

Herman, G.E. and El-Hodiri, H.M., 2002. The role of ZIC3 in vertebrate development. Cytogenet. Genome. Res. 99:229–235.

Huisman, E., Uylings, H.B. and Hoogland, P.V. 2004. A 100% increase of dopaminergic cells in the olfactory bulb may explain hyposmia in Parkinson’s disease. Mov. Disord.: official journal of the Movement Disorder Society. 19:687–692.

Hwang, D.Y., Ardayfio, P., Kang, U.J., Semina, E.V. and Kim, K.S. 2003. Selective loss of dopaminergic neurons in the substantia nigra of Pitx3-deficient aphakia mice. Mol. Brain. Res. 114:123–131.

Inoue, T., Ota, M., Ogawa, M., Mikoshiba, K. and Aruga, J. 2007. Zic1 and Zic3 regulate medial forebrain development through expansion of neuronal progenitors. J. Neurosci. 27:5461–5473.

Jessberger, S., Clark, R.E., Broadbent, N.J., Clemenson, G.D., Consiglio, A., Lie, D.C., Squire, L.R. and Gage, F.H. 2009. Dentate gyrus-specific knockdown of adult neurogenesis impairs spatial and object recognition memory in adult rats. Learn. Mem. 16:147–154.

Kadkhodaei, B., Ito, T., Joodmardi, E., Mattsson, B., Rouillard, C., Carta, M., Muramatsu, S.I., Sumi-Ichinose, C., Nomura, T., Metzger, D. and Chambon, P. 2009. Nurr1 is required for maintenance of maturing and adult midbrain dopamine neurons. J.Neurosci. 29:15923–15932.

Kitaguchi, T., Mizugishi, K., Hatayama, M., Aruga, J. and Mikoshiba, K. 2002. Xenopus Brachyury regulates mesodermal expression of Zic3, a gene controlling left–right asymmetry. Dev. Growth. Differ. 44:55–61.

Kitaguchi, T., Nagai, T., Nakata, K., Aruga, J. and Mikoshiba, K. 2000. Zic3 is involved in the left-right specification of the Xenopus embryo. Development, 127:4787–4795.

Klootwijk, R., Franke, B., van der Zee, C.E., de Boer, R.T., Wilms, W., Hol, F.A. and Mariman, E.C. 2000. A deletion encompassing Zic3 in bent tail, a mouse model for X-linked neural tube defects. Hum. Mol. Genet. 9:1615–1622.

Kohwi, M., Osumi, N., Rubenstein, J.L. and Alvarez-Buylla, A. 2005. Pax6 is required for making specific subpopulations of granule and periglomerular neurons in the olfactory bulb. J. Neurosci. 25: 6997–7003.

Kohwi, M., Petryniak, M.A., Long, J.E., Ekker, M., Obata, K., Yanagawa, Y., Rubenstein, J.L. and Alvarez-Buylla, A. 2007. A subpopulation of olfactory bulb GABAergic interneurons is derived from Emx1-and Dlx5/6-expressing progenitors. J. Neurosci. 27:6878–6891.

Koyabu, Y., Nakata, K., Mizugishi, K., Aruga, J. and Mikoshiba, K. 2001. Physical and functional interactions between Zic and Gli proteins. J. Biol. Chem. 276:6889–6892.

Kruzich, P.J. and Grandy, D.K. 2004. Dopamine D 2 receptors mediate two-odor discrimination and reversal learning in C57BL/6 mice. BMC neurosci. 5:1–10.

Lim, L.S., Hong, F.H., Kunarso, G. and Stanton, L.W. 2010. The pluripotency regulator Zic3 is a direct activator of the Nanog promoter in ESCs. Stem cells. 28:1961–1969.

Lim, L.S., Loh, Y.H., Zhang, W., Li, Y., Chen, X., Wang, Y., Bakre, M., Ng, H.H. and Stanton, L.W. 2007. Zic3 is required for maintenance of pluripotency in embryonic stem cells. Mol. Biol. Cell. 18:1348–1358.

Liu, N. and Baker, H. 1999. Activity-dependent Nurr1 and NGFI-B gene expression in adult mouse olfactory bulb. Neuroreport, 10:747–751.

Lledo, P.M., Merkle, F.T. and Alvarez-Buylla, A. 2008. Origin and function of olfactory bulb interneuron diversity. Trends Neurosci. 31:392–400.

Lois, C., Garcia-Verdugo, J.M. and Alvarez-Buylla, A. 1996. Chain migration of neuronal precursors. Science. 271:978–981.

Marchal, L., Luxardi, G., Thomé, V. and Kodjabachian, L. 2009. BMP inhibition initiates neural induction via FGF signaling and Zic genes. Proc. Nat. Acad. Sci. 106:17437–17442.

Mizugishi, K., Aruga, J., Nakata, K. and Mikoshiba, K. 2001. Molecular properties of Zic proteins as transcriptional regulators and their relationship to GLI proteins. J. Biol. Chem. 276:2180–2188.

Mizuseki, K., Kishi, M., Matsui, M., Nakanishi, S. and Sasai, Y. 1998. Xenopus Zic-related-1 and Sox-2, two factors induced by chordin, have distinct activities in the initiation of neural induction. Development. 125:579–587.

Morley JF, Cohen A, Silveira-Moriyama L, Lees AJ, Williams DR, Katzenschlager R, Hawkes C, Shtraks JP, Weintraub D, Doty RL, Duda JE. 2018. Optimizing olfactory testing for the diagnosis of Parkinson’s disease: item analysis of the university of Pennsylvania smell identification test. npj Parkinson’s Dis. 15;4(1):1–7.

Nagayama S, Homma R, Imamura F. Neuronal organization of olfactory bulb circuits. 2014. Front. neural circuits. 3;8:98.

Nakata, K., Koyabu, Y., Aruga, J. and Mikoshiba, K. 2000. A novel member of the Xenopus Zic family, Zic5, mediates neural crest development. Mech. Dev. 99:83–91.

Nakata, K., Nagai, T., Aruga, J. and Mikoshiba, K. 1997. Xenopus Zic3, a primary regulator both in neural and neural crest development. Proc.Nat. Acad. Sci. 94:11980–11985.

Parrish-Aungst, S., Shipley, M.T., Erdelyi, F., Szabo, G. and Puche, A.C. 2007. Quantitative analysis of neuronal diversity in the mouse olfactory bulb. J. Comp. Neurol. 501:825–836.

Purandare, S.M., Ware, S.M., Kwan, K.M., Gebbia, M., Bassi, M.T., Deng, J.M., Vogel, H., Behringer, R.R., Belmont, J.W. and Casey, B. 2002. A complex syndrome of left-right axis, central nervous system and axial skeleton defects in Zic3 mutant mice. 2293–2302.

Qin, S., Ware, S.M., Waclaw, R.R. and Campbell, K. 2017. Septal contributions to olfactory bulb interneuron diversity in the embryonic mouse telencephalon: role of the homeobox gene Gsx2. Neural. Dev. 12:1–14.

Remesal, L., Roger-Baynat, I., Chirivella, L., Maicas, M., Brocal-Ruiz, R., Pérez-Villalba, A., Cucarella, C., Casado, M. and Flames, N. 2020. PBX1 acts as terminal selector for olfactory bulb dopaminergic neurons. Development. 147:186841.

Rey NL, Wesson DW, Brundin P. The olfactory bulb as the entry site for prion-like propagation in neurodegenerative diseases. 2018. Neurobiol of dis. 1;109:226–48.

Saino-Saito, S., Cave, J.W., Akiba, Y., Sasaki, H., Goto, K., Kobayashi, K., Berlin, R. and Baker, H. 2007. ER81 and CaMKIV identify anatomically and phenotypically defined subsets of mouse olfactory bulb interneurons. J. Comp. Neurol. 502:485–496.

Sakurada, K., Ohshima-Sakurada, M., Palmer, T.D. and Gage, F.H. 1999. Nurr1, an orphan nuclear receptor, is a transcriptional activator of endogenous tyrosine hydroxylase in neural progenitor cells derived from the adult brain. Development 126:4017–4026.

Sasai, N., Mizuseki, K. and Sasai, Y. 2001. Requirement of FoxD3-class signaling for neural crest determination in Xenopus. 2525–2536.

Saucedo-Cardenas, O. and Conneely, O.M. 1996. Comparative distribution of NURR1 and NUR77 nuclear receptors in the mouse central nervous system. J. Mol. Neurosci. 7:51–63.

Smits, S.M. and Smidt, M.P. 2006. The role of Pitx3 in survival of midbrain dopaminergic neurons. Parkinson’s Disease. Relat. Disord. 57–60.

Sone, M., Morone, N., Nakamura, T., Tanaka, A., Okita, K., Woltjen, K., Nakagawa, M., Heuser, J.E., Yamada, Y., Yamanaka, S. and Yamamoto, T. 2017. Hybrid cellular metabolism coordinated by Zic3 and Esrrb synergistically enhances induction of naive pluripotency. Cell. Metabol. 25:1103–1117.

Specht, L.A., Pickel, V.M., Joh, T.H. and Reis, D.J. 1981. Light-microscopic immunocytochemical localization of tyrosine hydroxylase in prenatal rat brain. I. Early ontogeny. J. Comp. Neurol. 199:233–253.

Stenman, J., Toresson, H. and Campbell, K. 2003. Identification of two distinct progenitor populations in the lateral ganglionic eminence: implications for striatal and olfactory bulb neurogenesis. J. Neurosci. 23:167–174.

Tiveron, M.C., Beclin, C., Murgan, S., Wild, S., Angelova, A., Marc, J., Coré, N., de Chevigny, A., Herrera, E., Bosio, A. and Bertrand, V. 2017. Zic-proteins are repressors of dopaminergic forebrain fate in mice and C. elegans. J. Neurosci. 37:10611–10623.

Vassar, R., Chao, S.K., Sitcheran, R., Nun, J.M., Vosshall, L.B. and Axel, R. 1994. Topographic organization of sensory projections to the olfactory bulb. Cell. 79:981–991.

Wang, S. and Turner, E.E. 2010. Expression of dopamine pathway genes in the midbrain is independent of known ETS transcription factor activity. J. Neurosci. 30:9224–9227.

Winata, C.L., Kondrychyn, I., Kumar, V., Srinivasan, K.G., Orlov, Y., Ravishankar, A., Prabhakar, S., Stanton, L.W., Korzh, V. and Mathavan, S. 2013. Genome wide analysis reveals Zic3 interaction with distal regulatory elements of stage specific developmental genes in zebrafish. PLoS. Genet. 9:1003852.

Xiao, Q., Chen, S. and Le, W. 2014. Hyposmia: a possible biomarker of Parkinson’s disease. Neuroscience bulletin. 30:134–140.

Yang, S.H., Andrabi, M., Biss, R., Baker, S.M., Iqbal, M. and Sharrocks, A.D. 2019. ZIC3 controls the transition from naive to primed pluripotency. Cell Rep. 27:3215–3227.

Yoneshima, H., Yamasaki, S., Voelker, C.C.J., Molnar, Z., Christophe, E., Audinat, E., Takemoto, M., Nishiwaki, M., Tsuji, S., Fujita, I. and Yamamoto, N. 2006. Er81 is expressed in a subpopulation of layer 5 neurons in rodent and primate neocortices. Neurosci. 137:401–412.

Zhao, C., Deng, W. and Gage, F.H. 2008. Mechanisms and functional implications of adult neurogenesis. Cell. 132:645–660.

