## Supplementary File for "Zic3 enables bimodal regulation of tyrosine hydroxylase expression in dopaminergic neurons of olfactory bulb and midbrain"

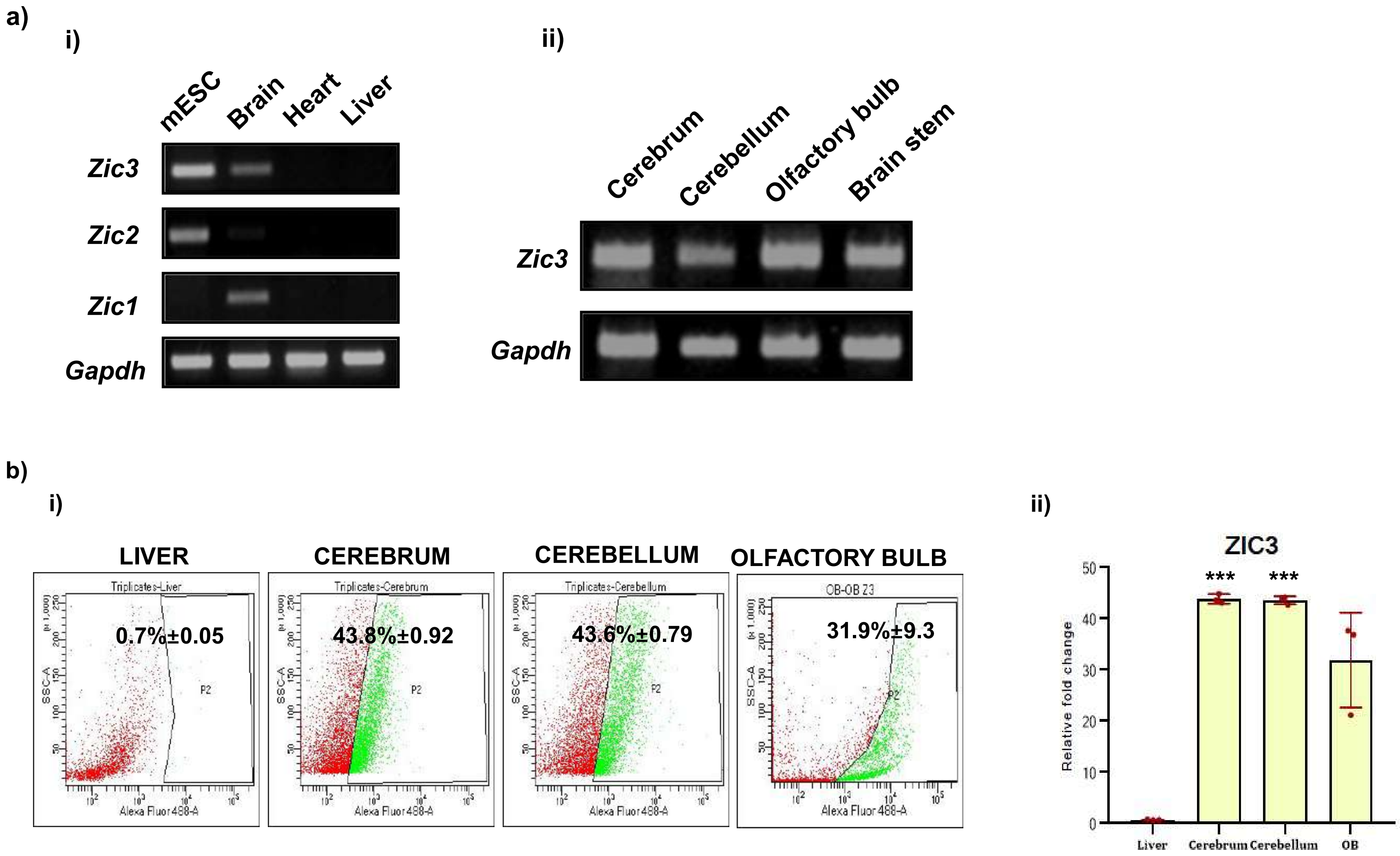

a)
i)
mESC
Brain
Heart
Liver
Zic3
Zic2
Zic1
Gapdh
ii)
Cerebrum
Cerebellum
Olfactory bulb
Brain stem
Zic3
Gapdh
b)
i)
LIVER
CEREBRUM
CEREBELLUM
OLFACTORY BULB
0.7%±0.05
43.8%±0.92
43.6%±0.79
31.9%±9.3
Alexa Fluor 488-A
SSC-A
ii)
ZIC3
Relative fold change
***
***
Liver
Cerebrum
Cerebellum
OB

**Figure S1: ZIC3 is expressed in various regions of brain including olfactory bulb**

(a) Transcript analysis of *Zic3* in various mouse tissues (i) and different regions of brain (ii), (b) flow cytometry analysis (i) and quantification (ii) of ZIC3 in different tissues of mouse. Scale bar represents 100 µm. Mean+/-SE of biological triplicates, *** $p \leq 0.001$.

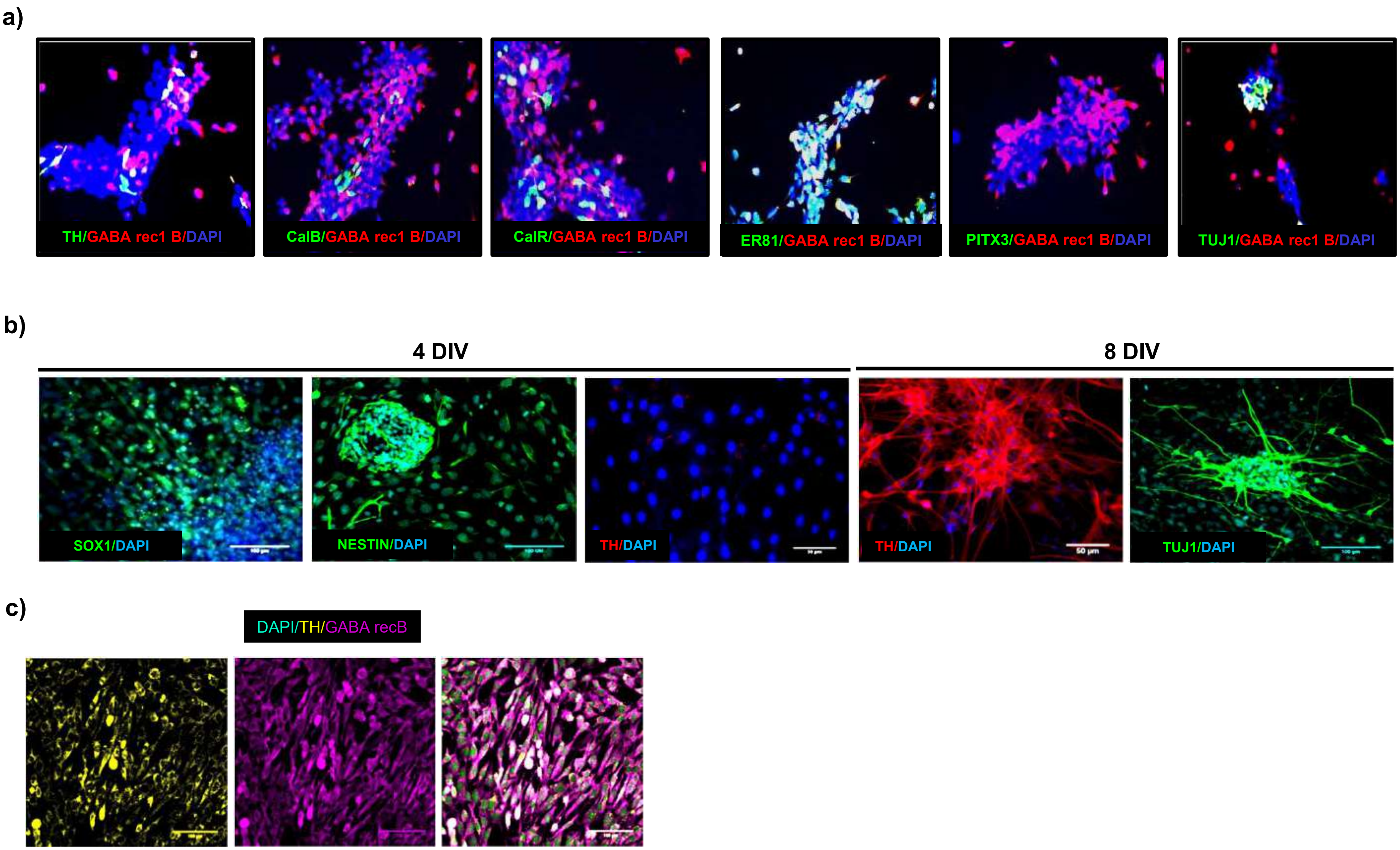

a)
TH/GABA rec1 B/DAPI
CalB/GABA rec1 B/DAPI
CalR/GABA rec1 B/DAPI
ER81/GABA rec1 B/DAPI
PITX3/GABA rec1 B/DAPI
TUJ1/GABA rec1 B/DAPI
b)
4 DIV
8 DIV
SOX1/DAPI
NESTIN/DAPI
TH/DAPI
TH/DAPI
50 µm
TUJ1/DAPI
c)
DAPI/TH/GABA recB

**Figure S2: Characterization of OB primary neurons:** (a) Expression of TH, CalB, CalR, ER81, PITX3 and TUJ1 in OB primary cells isolated from embryonic mouse OB, (b) immunofluorescence staining of SOX1, NESTIN in OB primary cells after 4 DIV; TH and TUJ1 on 8th DIV in OB DA neuron differentiation, (c) co-expression of TH and GABA rec1 B in differentiated OB DA primary neurons. Scale bar represents 100 μm.

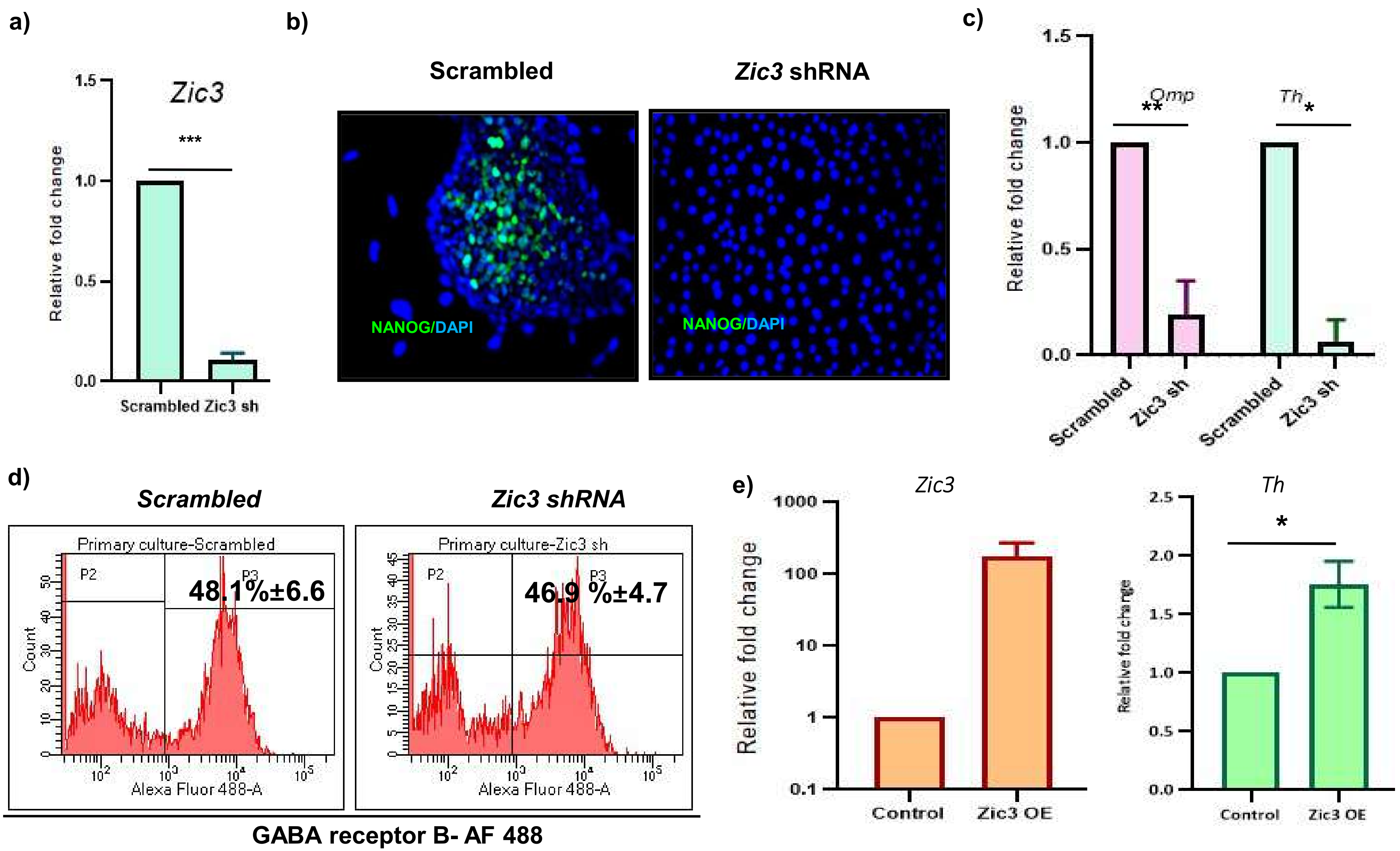

a)
Zic3
***
1.5
1.0
0.5
0.0
Relative fold change
Scrambled Zic3 sh
b)
Scrambled
Zic3 shRNA
NANOG/DAPI
NANOG/DAPI
c)
1.5
1.0
0.5
0.0
Relative fold change
Omp
**
Th
*
Scrambled
Zic3 sh
Scrambled
Zic3 sh
d)
Scrambled
Zic3 shRNA
Primary culture-Scrambled
P2
P3
48.1%±6.6
Count
Alexa Fluor 488-A
Primary culture-Zic3 sh
P2
P3
46.9 %±4.7
Count
Alexa Fluor 488-A
GABA receptor B- AF 488
e)
Zic3
1000
100
10
1
0.1
Relative fold change
Control
Zic3 OE
Th
*
2.5
2.0
1.5
1.0
0.5
0.0
Relative fold change
Control
Zic3 OE

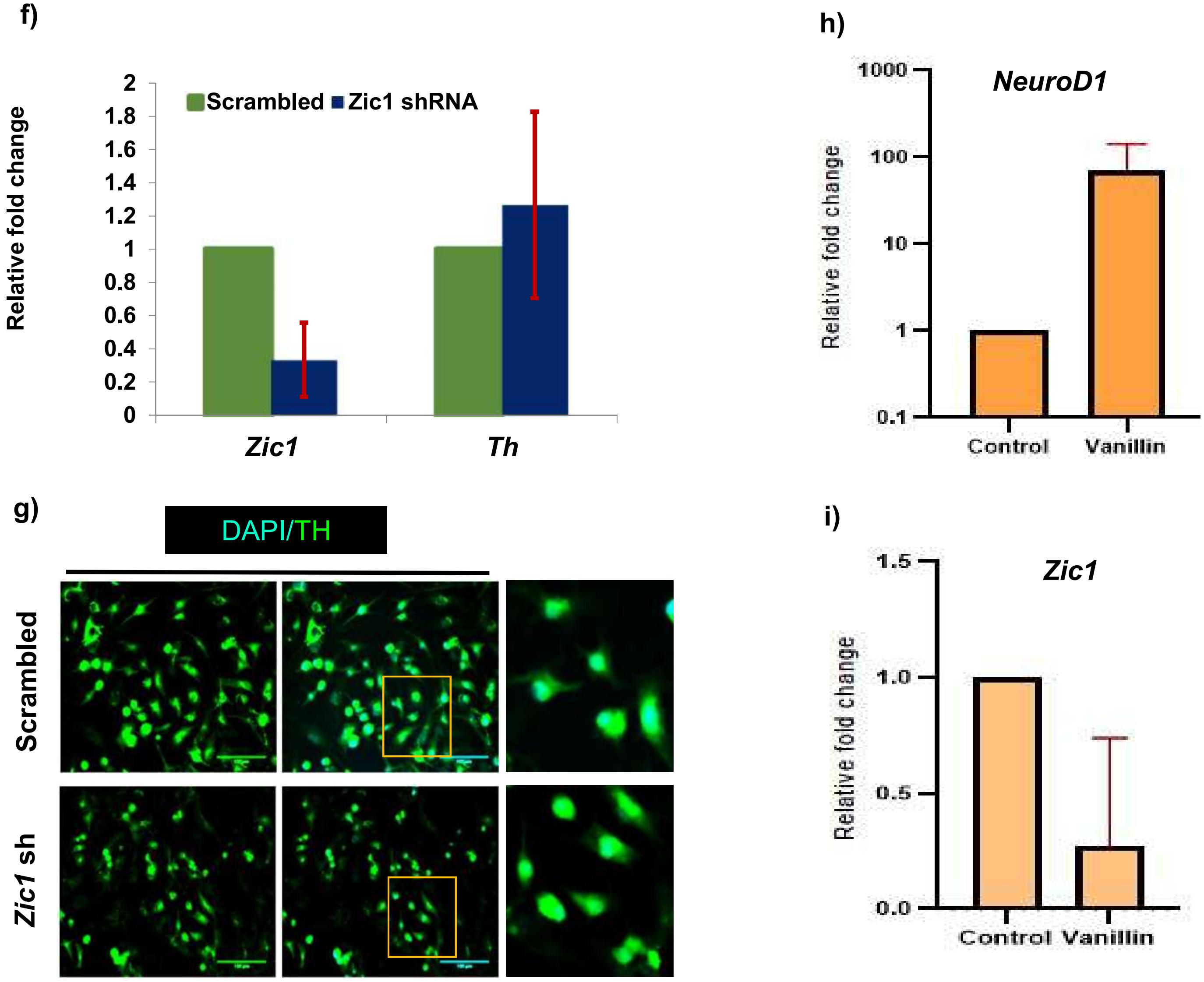
f)
Relative fold change
2
1.8
1.6
1.4
1.2
1
0.8
0.6
0.4
0.2
0
Scrambled
Zic1 shRNA
Zic1
Th
g)
DAPI/TH
Scrambled
Zic1 sh
h)
NeuroD1
Relative fold change
1000
100
10
1
0.1
Control
Vanillin
i)
Zic1
Relative fold change
1.5
1.0
0.5
0.0
Control
Vanillin

**Figure S3: *Zic3* is essential for and facilitates TH expression in OB primary DAergic neurons:** (a) Transcript analysis showing knockdown efficiency of *Zic3* shRNA in mESCs, (b) NANOG immunofluorescence showing functional effect of *Zic3* inhibition on mESC pluripotency, (c) Transcript analysis of *Th* and *Omp* expression in OB DA primary neurons transduced with *Zic3 shRNA*, (d) flow cytometric analysis of GABA rec1B expression showing no modulation upon *Zic3* shRNA in OB DA neurons, (e) mRNA analysis of *Zic3* and *Th* in cells over-expressing *Zic3*, transcript levels (f) and protein levels (g) of TH in OB DA neurons with *Zic1* shRNA, (h) Transcript analysis of NeuroD1 and (i) *Zic1* mRNA during vanillin treatment in primary OB . Mean+/-SE of biological triplicates, * $p \leq 0.05$, ** $p \leq 0.01$.

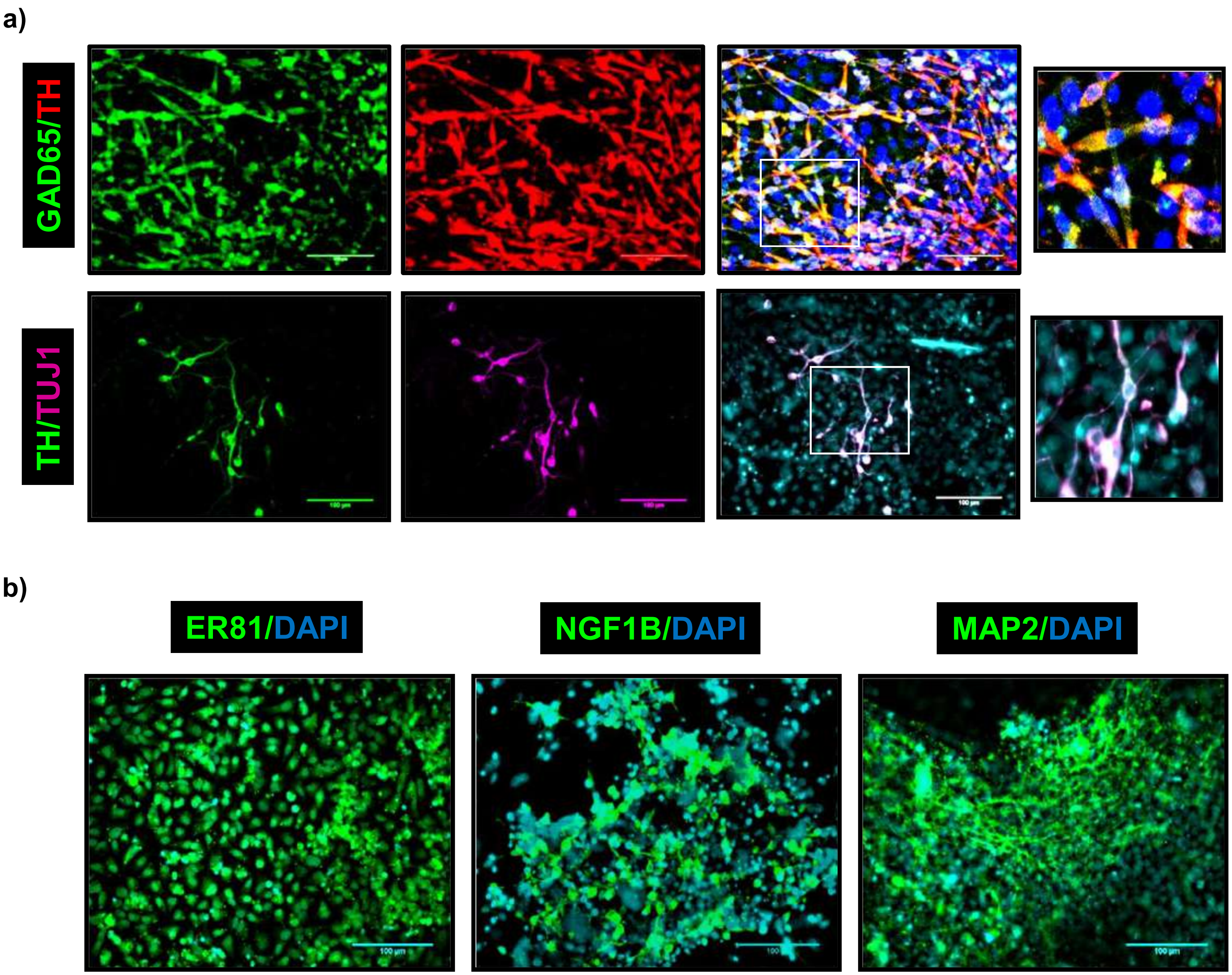

a)
GAD65/TH
TH/TUJ1
b)
ER81/DAPI
NGF1B/DAPI
MAP2/DAPI

**Figure S4: mESCs efficiently differentiate to OB DA like neurons *in vitro*:**
(a) Co-expression analysis of TH with GAD65 and TUJ1, b) staining for ER81, NGF1B and MAP2 in OB DA like neurons differentiated from mESCs. Scale bar represents 100 μm.

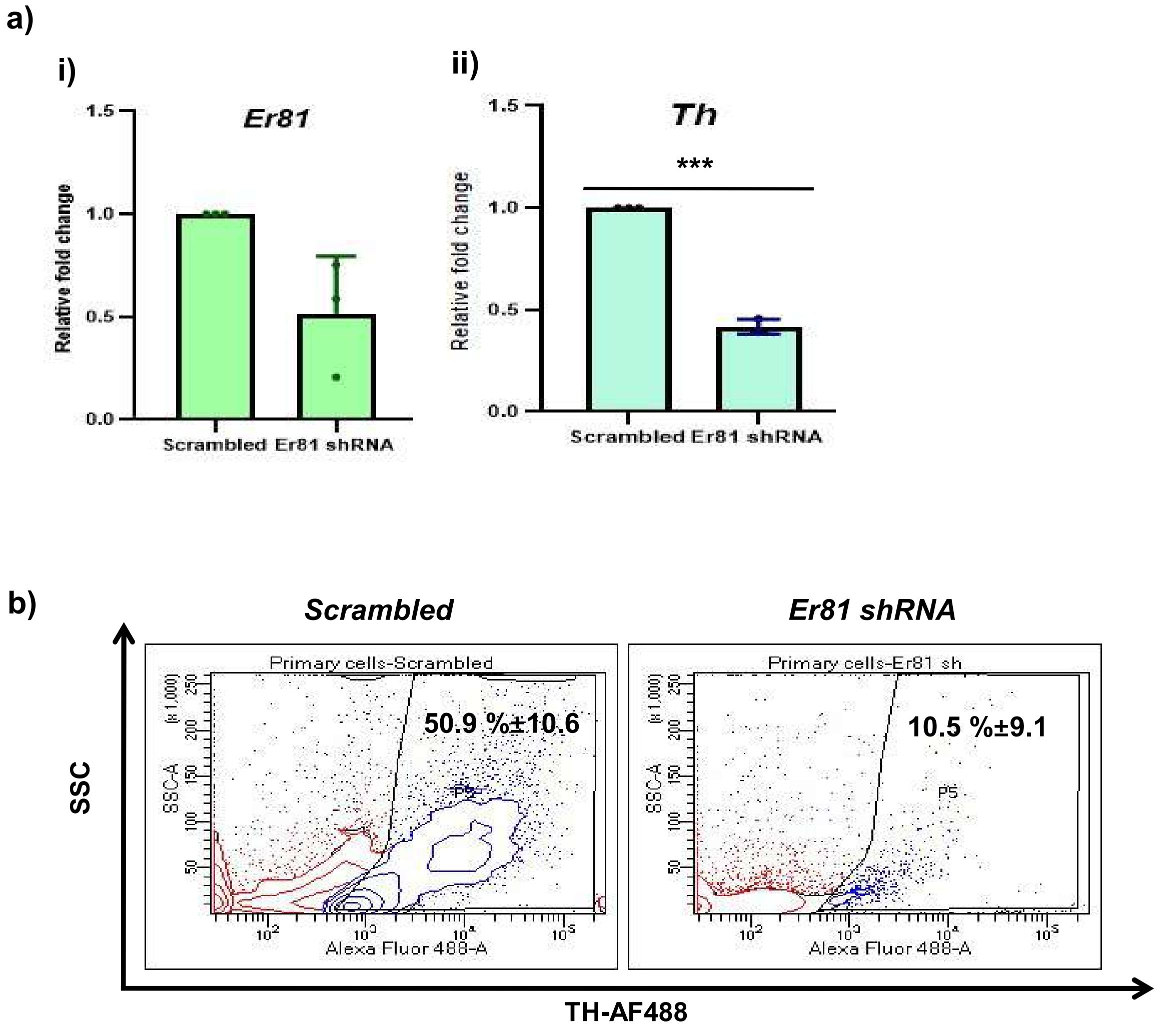

a)
i)
Er81
Relative fold change
1.5
1.0
0.5
0.0
Scrambled
Er81 shRNA
ii)
Th
***
Relative fold change
1.5
1.0
0.5
0.0
Scrambled
Er81 shRNA
b)
Scrambled
Er81 shRNA
Primary cells-Scrambled
Primary cells-Er81 sh
50.9 %±10.6
10.5 %±9.1
P5
SSC-A
Alexa Fluor 488-A
SSC
TH-AF488

**Figure S5: *Er81* is essential for TH expression in OB DA like neurons**

Transcript analysis of *Er81* (a. i) and *Th* (a. ii) in cells harboring *Er81* shRNA construct (b) flow cytometric analysis of TH upon *Er81* knockdown in OB DA primary neurons. Mean+/-SE of biological triplicates, *** p≤0.001.

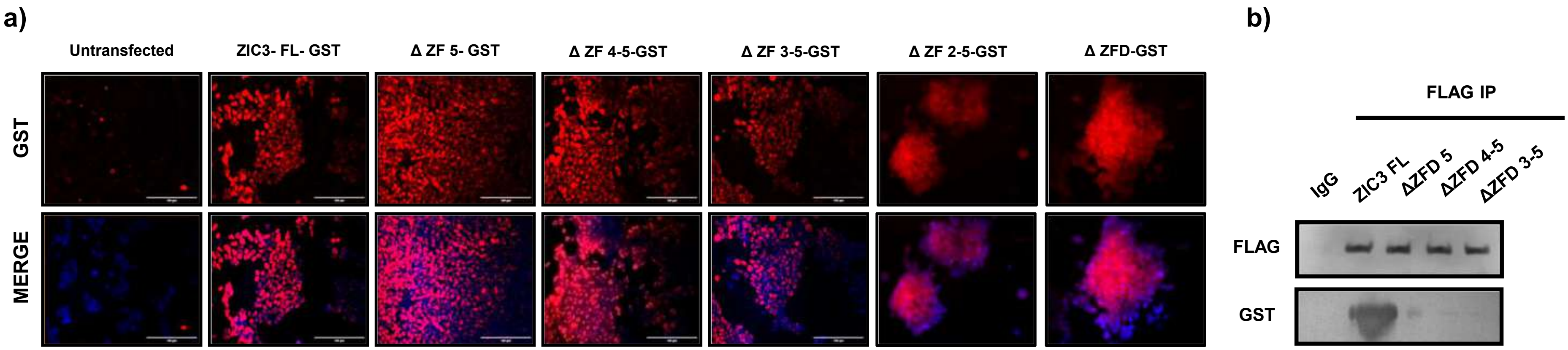
a)
Untransfected
ZIC3- FL- GST
Δ ZF 5- GST
Δ ZF 4-5-GST
Δ ZF 3-5-GST
Δ ZF 2-5-GST
Δ ZFD-GST
GST
MERGE
b)
FLAG IP
IgG
ZIC3 FL
ΔZFD 5
ΔZFD 4-5
ΔZFD 3-5
FLAG
GST

**Figure S6: Deletion of ZIC3 zinc fingers leads to cytoplasmic accumulation of ZIC3 and hinders its interaction with ER81**
(a) immunofluorescence showing sub-cellular localization of ZIC3 GST FL and zinc finger mutants, (b) western blot to show lack of interaction between zinc finger mutants of ZIC3 and ER81. Scale bar represents 100 μm.

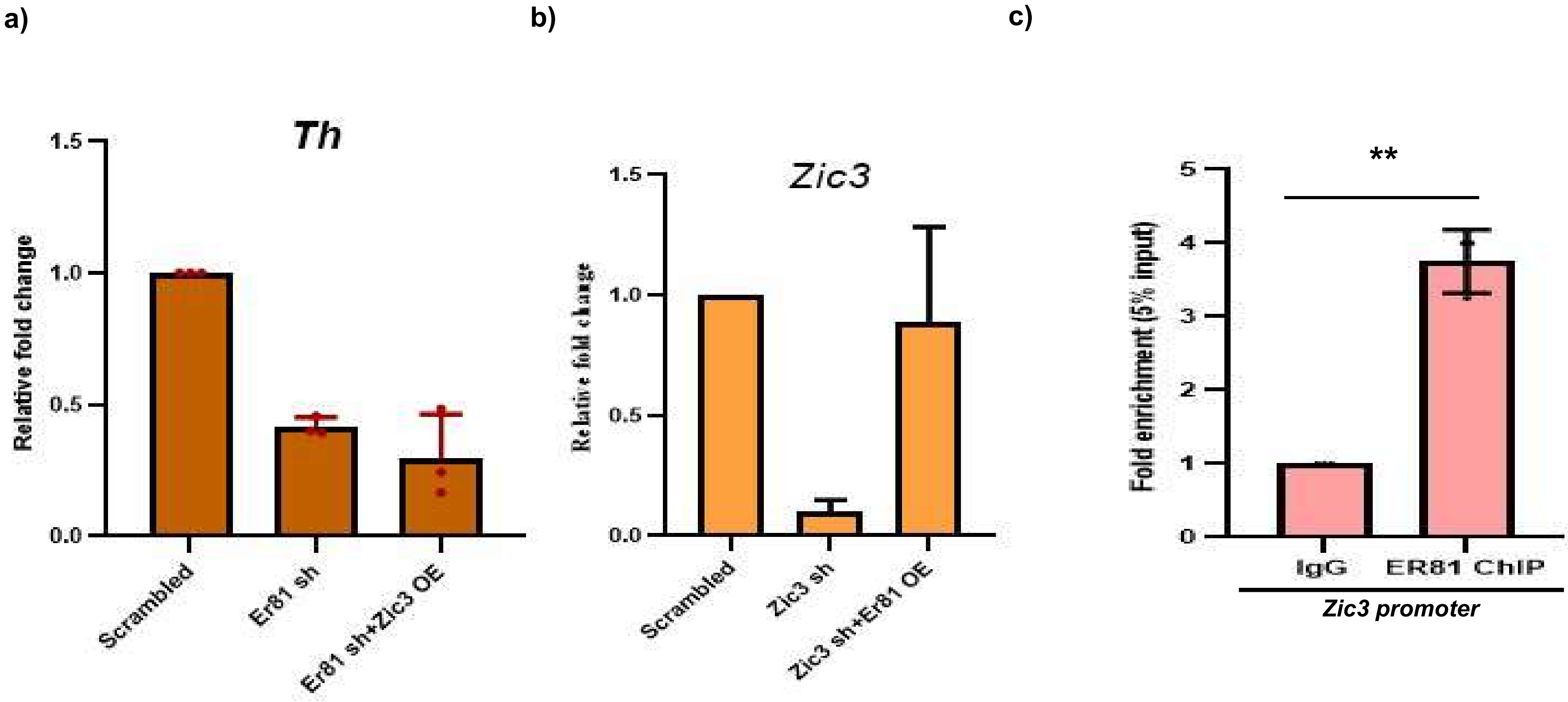
a)
Th
1.5
1.0
0.5
0.0
Relative fold change
Scrambled
Er81 sh
Er81 sh+Zic3 OE
b)
Zic3
1.5
1.0
0.5
0.0
Relative fold change
Scrambled
Zic3 sh
Zic3 sh+Er81 OE
c)
**
5
4
3
2
1
0
Fold enrichment (5% input)
IgG
ER81 ChIP
Zic3 promoter

**Figure S7: ZIC3 fails to rescue *Er81* knockdown mediated inhibition of *Th* expression**

(a) effect of *Zic3* over-expression on *Th* mRNA levels in OB DA neurons with *Er81* loss of function, (b) Expression of *Zic3* mRNA in cells containing *Zic3* shRNA and *Er81* over-expression plasmid constructs, (c) ChIP PCR with ER81 antibody showing binding of ER81 to *Zic3* promoter. Mean+/-SE of biological triplicates, ** $p \leq 0.01$.

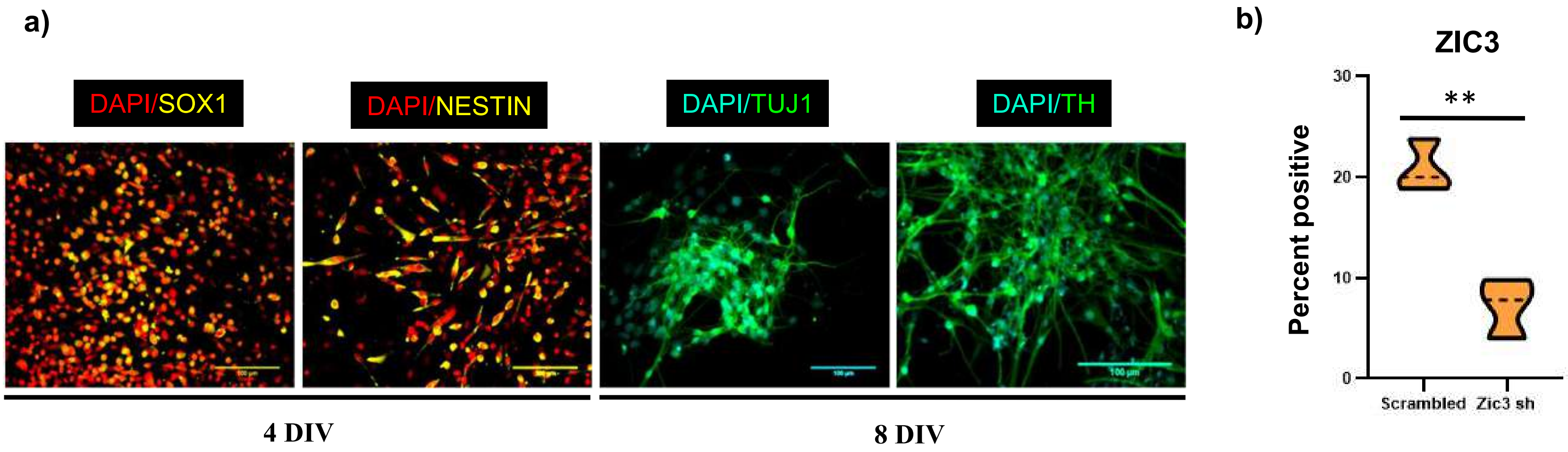

a)
DAPI/SOX1
DAPI/NESTIN
DAPI/TUJ1
DAPI/TH
4 DIV
8 DIV
b)
ZIC3
Percent positive
30
20
10
0
**
Scrambled
Zic3 sh

**Figure S8: Derivation of MB DA like neurons from primary neurospheres**

(a) Immunofluorescence showing the expression of neural progenitor markers SOX1 and NESTIN in MB primary cells 4 DIV and matured neural markers TUJ1 and DA marker TH 8 DIV, (b) knockdown efficiency of ZIC3 in MB DA neurons as shown by quantification of flow cytometric staining. Scale bar represents 100 µm. Mean+/-SE of biological triplicates, ** $p \leq 0.01$.

***Supplementary table 1***

| Primer name | Sequence | Amplicon size (bp) |
|---|---|---|
| *mZic3* | Forward primer:AAGATTTTTGCCCGCTCTG<br>Reverse primer:TATAGGGCTTGTCCGAGGTG | 157 |
| *mGapdh* | Forward primer: ACCACAGTCCATGCCATCAC<br>Reverse primer: TCCACCACCCTGTTGCTGTA | 425 |
| *mOct 4* | Forward primer: GAGGAGTCCAGGACATGAA<br>Reverse primer: AGATGGTGGTCTGGCTGAAC | 153 |
| *mMap2* | Forward primer: TCAGGAGACAGGGAGGAGAA<br>Reverse primer: GTGTGGAGGTGCCACTTTTT | 112 |
| *mZic1* | Forward primer: GCCCTTCAAAGCCAAATACA<br>Reverse primer: TTGCAAAGGTAGGGCTTGTC | 252 |
| *mTh* | Forward primer:AGGAGAGGGATGGAAATGCT<br>Reverse primer: GCGCACAAAGTACTCCAGGT | 179 |
| *mGad65* | Forward primer:AGATCGCCCCTGTATTTGTG<br>Reverse primer:GCATGGCATACATGTTGGAG | 132 |
| *mEr81* | Forward primer:TTCAGAACTCGGGTCTGCTT<br>Reverse primer:TGAGCTGTGTTTGGAGATGC | 183 |
| *mNurr1* | Forward primer:AGTCTGATCAGTGCCCTCGT<br>Reverse primer:GATCTCCATAGAGCCGGTCA | 162 |
| *mEngrailed* | Forward primer:GACTCTTCAGGCATCCAAGC<br>Reverse primer:GGGTCATCCAGTGCTGCTAT | 258 |
| *mAadc* | Forward primer:CTGATGTGGAGCCTGGCTAT<br>Reverse primer:GAGAAACCAATGCAGCCAAT | 220 |
| *mVmat2* | Forward primer:CCTCTGCTGGTGGTGCTATT<br>Reverse primer:CTTAATGGGGCAGTTGTGGT | 164 |
| *mGch1* | Forward primer:CAAGCAAGTCCTTGGTCTCA<br>Reverse primer:GAGGAACTCCTCCCGAGTCT | 259 |
| *mNgf1b* | Forward primer: TTCTGCTCAGGCCTGGTACT<br>Reverse primer: AATGCGATTCTGCAGCTCTT | 204 |
| *mPitx3* | Forward primer: GCAACTGGCCGCCCAAGG<br>Reverse primer: AGGCCCCACGTTCACCGA | 84 |
| m*Th* promoter luciferase | Forward primer: GAGGCCTCTTGGGATT<br>Reverse primer: CTGGTGGTCCCGAGTT | 2559 |
| m*Zic3* EBS | Forward primer: AAGCTGACAGGATCCAAAC<br>Reverse primer: TAAATCATGCAAATGAATTC | 180 |
| *Th* EBS (+7 to -99 bp) | Forward primer: TGGATGCAATTAGATCTAATGGGACG<br>Reverse primer: TGGGCATAGTGCAAGCTGGTGGTCCC | 106 |
| *Th* (-49 to -245 bp) | Forward primer: GGATCTTTGTGTAAGTGG<br>Reverse primer: AGTTAAGAGTATCCTGAAC | 196 |
| *Th* (-246 to -434 bp) | Forward primer: AAAGCAGAGGTCTGTCCC<br>Reverse primer: GTCCTATGAGACACAGAA | 189 |
| *Th* (-435 to -629 bp) | Forward primer: CTGCCTGAGGACCCAGCC<br>Reverse primer: GTTCATGTTAGGAAGGC | 195 |
| m*Pitx3*ChIP | Forward primer: CCAAATCCTGCTTTCTCC<br>Reverse primer: CTATTCCAGTCCTCGTGC | 185 |

| | | |
|---|---|---|
| m*Zic3* FL GST | Forward primer: ATGACGATGCTCCTGGAC<br>Reverse primer: GACGTACCATTCGTT | 3366 |
| m*Zic3* ΔZF5 GST | Forward primer: ATGACGATGCTCCTGGAC<br>Reverse primer: CGAGGTGTGCACATGCATG | 2273 |
| m*Zic3* Δ ZF4-5 GST | Forward primer: ATGACGATGCTCCTGGAC<br>Reverse primer: ACCTGTATGGGTCCTCTTG | 1059 |
| m*Zic3* ΔZF3-5 GST | Forward primer: ATGACGATGCTCCTGGAC<br>Reverse primer: GCCAGTGTGCACTCGGATA | 969 |
| m*Zic3* Δ ZF2-5 GST | Forward primer: ATGACGATGCTCCTGGAC<br>Reverse primer: GTTGTTCTGCTCCGGGCC | 879 |
| mHA *Zic3* ZFD | Forward primer: GAGCTGTCCTGTAAGTGG<br>Reverse primer: TCAGACGTACCATTCGTTA | 2625 |
| m*ER81* FLAG | Forward primer: ATGGATGGATTTTA<br>Reverse primer: TTAGTACACGTATC | 1455 |
| m*Zic3* promoter GFP | Forward primer: ACCAGGGGGAAGAGTGGTG<br>Reverse primer: GGGGAACCACGGGGCCAG | 1451 |

***Supplementary table 2***

| Antibodies | Source | Identifier |
|---|---|---|
| Goat anti ZIC3 | Santa Cruz | #SC-28156 |
| Rabbit anti ZIC3 | ABclonal | #PA5-97216 |
| Rabbit anti TH | ABclonal | #A12756 |
| Mouse anti β-ACTIN | Santa Cruz | #SC-47778 |
| Rabbit anti NANOG | Millipore | #AB9220 |
| Chicken anti MAP2 | Abcam | #ab5392 |
| Rabbit anti TUJ1 | Abcam | #ab18207 |
| Mouse anti GAD65 | Abcam | #ab26113 |
| Rabbit anti ER81 | Abzyme | #AB-PA002391 |
| Rabbit anti NGF1B | CusaBio | #CSB-PA003493 |
| Mouse anti NESTIN | BD Pharmingen | #556309 |
| Mouse anti GABA B receptor 1 | Abcam | #ab55051 |
| Rabbit anti CALBINDIN | ABclonal | #A0802 |
| Rabbit anti CALRETININ | ABclonal | #A9807 |
| Rabbit anti PITX3 | ABclonal | #A19261 |
| Mouse anti DDDDK-tag (Anti-FLAG) | ABclonal | #AE005 |
| Rabbit anti HA-tag | ABclonal | #AE036 |
| Rabbit anti GST | ABclonal | #AE006 |
| Rabbit anti SOX1 | ABcam | #ab109290 |
| Anti PSA-NCAM | Miltenyi Biotec | #120-004-201 |

| | | |
|---|---|---|
| Anti A2B5 | Miltenyi Biotec | #120-005-044 |
| Chicken anti GFP | Aves labs | #GFP-1020 |
| Rabbit anti TH | EMD Millipore | #AB152 |
| Rabbit anti mouse FITC | Sigma Aldrich | #AP160F |
| Goat anti mouse AF594 | ThermoFisher Scientific | #11005 |
| Donkey anti rabbit AF594 | ThermoFisher Scientific | #21207 |
| Donkey anti rabbit AF488 | ThermoFisher Scientific | #21206 |
| Goat anti chicken AF488 | Invitrogen | #A-11039 |
| Rabbit anti mouse HRP | Southern Biotech | #6170-05 |
| Goat anti rabbit HRP | Abcam | GR209629-6 |
| Chicken anti goat FITC | Sigma Aldrich | #AP163F |
